# ABCG transporter knockouts alter integument and eye pigmentation with gene-specific effects on viability in butterflies and moths

**DOI:** 10.64898/2026.02.24.707689

**Authors:** Donya N. Shodja, Joseph Yan, Luca Livraghi, Amika Yamada, Arnaud Martin, Anyi Mazo-Vargas

## Abstract

Insects possess remarkable diversity in pigmentation across life stages, generated from a limited number of pigment precursors. ABCG transporters play central roles in pigmentation by mediating the intracellular transport of these precursors, yet their functions outside a few model systems remain poorly characterized. Here, we investigate the roles of the ABCG transporter genes *white (w)*, *scarlet (st)*, and *oily kinshiryu (ok)* across five lepidopteran species representing both butterflies and moths. CRISPR-Cas9 knockouts reveal conserved, tissue-specific functions of ABCG transporters throughout development, affecting larval and pupal integuments as well as adult eyes, but not scale coloration. Notably, our data indicate that both ommochrome and uric acid deposition underlie the coloration and opacity of Lepidoptera larvae and pupae, with the relative contributions of these pathways differing across species and stages. We also reveal that the vivid orange coloration of Gulf fritillary caterpillars arises from the interplay of ommochromes and urate granules, providing a clear example of how these pathways combine to produce diverse integument pigmentation. Loss of *white* was deleterious in four species examined, whereas *scarlet* knockouts produced viable individuals with easily detectable pigmentation phenotypes, identifying *scarlet* mutant backgrounds as a practical genetic marker for functional studies. Together, these findings show that conserved ABCG transporter complexes are differentially deployed across tissues and life stages, providing a mechanistic basis for the evolutionary diversification of pigmentation in Lepidoptera and enabling expanded functional genomics and transgenesis in emerging model species.

**Research Highlights:** CRISPR-Cas9 disruption of ABCG transporters across Lepidoptera reveals conserved, tissue-specific roles in pigmentation. Interactions between ommochrome and uric acid deposition generate stage-specific coloration in larval and pupal integument.

## INTRODUCTION

Insect pigmentation is a diverse and rapidly evolving trait that spans a wide range of tissues, and life stages, serving critical ecological and physiological roles throughout the life cycle. Pigmentation strategies include background matching, disruptive coloration, and mimicry, all of which are essential for survival in these often relatively immobile and vulnerable stages (Lindstedt et al., 2020). Coloration also serves physiological roles such as photoprotection (Hu et al., 2013), thermoregulation (Lahondère, 2023), and excretion via tissue storage of waste products (Ferré, 2024; Fujii & Banno, 2019). Insect colors arise from the synthesis or uptake of pigment molecules such as brown-black melanins, yellow-red-brown ommochromes, yellow-red-white pteridines, blue-green biliverdins, yellow carotenoids, and white uric acid, as well as from physical structures that generate structural colors (Futahashi & Osanai-Futahashi, 2021).

ABC transporters (ATP-binding cassette transporters) comprise a large, evolutionarily conserved protein family present across all domains of life that function as dimers to transport a wide range of substrates across cellular membranes (Holland, 2011; Srikant, 2020). Among them, White (*w*), Scarlet (*st*), Brown (*br*), and Oily kinshiryu (*ok*) are closely related transporters of the ABCG subfamily involved in pigmentation across insects (Mocchetti et al., 2025; Reding & Pick, 2020; Wang et al., 2013). These proteins assemble as heterodimers at the membranes of intracellular pigment granules, where they control the trafficking of pigment precursors that ultimately underlie the pigmentation of insect eyes, integuments, and other tissues (Figon & Casas, 2019; Mackenzie et al., 2000). Because of their combinatorial effects on specific types of pigments, mutants for these color determinants provide a powerful window into how distinct transporter heterodimers generate diverse pigmentation and physiological outcomes. Eye coloration defects in mutants for White (*w*), Scarlet (*st*), and Brown (*br*) have been extensively studied in *Drosophila melanogaster*. White acts as a shared subunit that heterodimerizes with Scarlet to transport tryptophan-derived ommochrome precursors, and with Brown to transport guanine-derived pteridine precursors (Ewart & Howells, 1998; Mackenzie et al., 2000). This division of labor underlies the canonical *Drosophila* eye-color phenotypes: loss of both pigment classes in *white* mutants yields white eyes, selective loss of reddish pteridines in *brown* mutants leaves ommochromes intact, and selective loss of brown ommochromes in *scarlet* mutants leaves pteridines intact. The transporter Ok, which is absent from *Drosophila* but otherwise widespread across insects (Mocchetti et al., 2025; Wang et al., 2013), partners with White to transport uric acid, a purine derivative, into urate granules in several lineages of Lepidoptera, resulting in white and opaque pigmentation of caterpillar integuments (Fujii & Banno, 2019; J. Lee et al., 2018; Ninomiya et al., 2006). Evolutionary comparisons of the roles of these transporters can offer a paradigm to understand how ABC complexes specialize, and more specifically here, how their recruitment has primed the diversification of color patterns seen in insects.

While the mechanisms of pigmentation are well studied in adult butterflies and moths (Lepidoptera), our knowledge of the genetic and molecular basis of pigmentation in their caterpillars and pupae is largely incomplete. To date, work mostly focused on the Bombycoidea and Papilionidae lineages has linked black larval color to melanins (Futahashi & Osanai-Futahashi, 2021), blue-green larval color to biliverdin and carotenoids (Barbier, 1981; Saito, 1998; Yoda et al., 2020), and white to uric acid (Fujii & Banno, 2019; J. Lee et al., 2018; Lhonoré et al., 1980; Ninomiya et al., 2006). These principles are likely to apply to other lepidopteran larvae and pupae that use camouflage strategies for survival. On the other hand, these pathways may be insufficient to explain the full gamut of colors observed across this enormous order, especially in lepidopteran caterpillars that are chemically defended and use vivid colors to signal their unpalatability to predators (Berenbaum, 1995). Testing the function of ABCG transporters may provide new insights into this phenomenon.

Of note, eye-color mutants such as *white* are often used as experimental tools in insect genetics, for example as genetic markers in crosses and transformation assays (Kane et al., 2017; Roseman et al., 1993, 1993), or by improving the detection of transgenesis markers that rely on eye fluorescence (Stern et al., 2017). However, *white* null mutations are lethal in several insect species (Khan et al., 2017; Lima et al., 2024; Reding & Pick, 2020), or can introduce artifacts in studies of behavior and physiology when they are viable (Ferreiro et al., 2018; Rickle, Sudhakar, et al., 2025). Thus, the functional study of alternative ABCG transporter genes may be useful for enabling functional genomics approaches across new laboratory insect model systems.

Here, we compare the function of the ABCG transporter genes *white*, *scarlet*, and *ok* across five lepidopteran species including three butterflies, the painted lady butterfly *Vanessa cardui,* the buckeye butterfly *Junonia coenia,* the gulf fritillary *Agraulis incarnata*, and two moths, the pantry moth *Plodia interpunctella* and the velvetbean caterpillar *Anticarsia gemmatalis*. CRISPR knockouts of these transporters showed distinct effects on the pigmentation of eggs, caterpillars, pupae, and adult eyes, notably supporting a role of ommochromes and uric acid in the pigmentation of larval integuments. While *white* loss-of-function proved deleterious in 4 out of 5 tested species, *scarlet* knockouts produced fully viable and easily screenable phenotypes, establishing *scarlet* knockouts as a more reliable genetic background for building transgenic lines.

## MATERIAL AND METHODS

### Insect rearing and husbandry

*J. coenia* and *V. cardui* colonies were respectively maintained at 28°C and 25°C in growth chambers with 60–70% relative humidity and a 14:10 light:dark photoperiod. Egg collection and larval rearing on artificial diets, and microinjections followed previously described methods (Tendolkar et al., 2021; Thulluru et al., 2022). *V. cardui* larvae were purchased from Carolina Biological Supply. *J. coenia* larvae were provided by Robert D. Reed from a colony originally established by Fred H. Nijhout. *Agraulis incarnata* colonies were maintained at 28°C in growth chambers with 60–70% relative humidity, under a 16:8 light:dark photoperiod, as well as in a greenhouse enclosure. Larvae were reared on *Passiflora biflora* or *Passiflora incarnata* plants. Egg collection and microinjections followed previously described methods (Hanly et al., 2023; Mazo-Vargas et al., 2022). Cultures of *P. interpunctella* used a wild-type laboratory strain (*bFog*) and were reared following previously published procedures (Heryanto et al., 2025; Heryanto, Hanly, et al., 2022; Heryanto, Mazo-Vargas, et al., 2022; Shodja et al., 2025). Briefly, larvae were reared at 28°C with 60-70% relative humidity and a 14:10 h light:dark cycle until adult emergence; adults were transferred to an oviposition jar following CO_2_ narcosis, which also induces mass egg laying by females. A first batch of eggs collected after 12 minutes was discarded in order to remove pre-fertilized eggs. For stock maintenance, synchronized eggs are collected in an aluminum cup, and 10-13 mg of eggs are transferred to a rearing container containing 45-50 g of compacted wheat bran diet, containing (w/w) 44% coarse wheat bran, 29% glycerol, 9% dextrose, 5% Brewer’s yeast, 10% water, 3% canola oil, 0.35% methylparaben, and 0.35% ascorbic acid. *A. gemmatalis* larvae were commercially obtained from Benzon Research and reared at 28°C and 60–70% relative humidity under a 14:10 light:dark photoperiod. Pupae were sorted by sex and adults were mixed together in a transparent Reptile Feeding Box (32 x 22 x 15 cm) for at least 12 hrs including cotton feeders with a 50% Gatorade solution (commercial drink diluted with an equal volume of water). For egg collection, stems of soybean (*Glycine max*) were put in floral water tubes filled with a water-based gel (the content of gel packs for cold shipping), and added to the container during the scotophase. Eggs were collected, transferred to a Vacuum Filtration Assembly Filter Kit, washed with 5% Benzalkonium Chloride and rinsed with distilled water, before incubation at 28°C. Cuttings of *Lablab purpureus* were provided in an insect proof containers after 72 hrs, and refreshed every 1-2 days for the first 5 days, before transfer of the larvae to cups containing the Southland Multiple Species diet. This larval diet was prepared with ∼90% of the water content recommended by the manufacturer (*e.g.* 850 mL instead of 930 mL) to reduce humidity in the feeding cups.

### Reagents for CRISPR mutagenesis of *white, scarlet* and *ok*

Cas9 short-guide RNA (sgRNA) targets were designed against the coding sequence of each target gene as listed in **Supporting Information Table S1,** using the reference genome of each species in NCBI (*V. cardui*, GCF_905220365.1; *P. interpunctella*, GCF_027563975.2 GCA_022985095.1 and GCA_001368715.1; *A. gemmatalis*, GCF_050436995.1; *A. incarnata* (listed as *Dione vanillae*), GCA_037178615.1), except for *J. coenia* (available at https://lepbase.cog.sanger.ac.uk/#archive/v4/sequence/). Genomes are listed in the CRISPOR database (https://crispor.gi.ucsc.edu/) for gRNA design (Concordet & Haeussler, 2018). Synthetic sgRNAs were provided by Synthego and Integrated DNA Technologies (IDT) and resuspended in Low TE buffer (10 mM Tris-HCl, 0.1mM EDTA, pH. 8.0) in 2.5 μL aliquots (500 ng/μL) and stored at - 70°C. Cas9-2xNLS recombinant protein (QB3/Macrolabs, UC Berkeley) was resuspended at 1,000 ng/μL in *Bombyx* 2X injection buffer (0.15 mM NaH₂PO₄, 0.85 mM Na₂HPO₄, and 10 mM KCl, pH 7.2–7.4) containing 0.05% cell-culture grade Phenol Red, and stored at -70°C in 2.5 μL aliquots. CRISPR- injection mixes consisted of freshly prepared mixes of equal volumes of aliquoted Cas9:sgRNA to a final concentration of 500:250 ng/μL.

### Microinjection reagents for *pBac* transgenesis

The donor plasmids *pBac{3XP3::EYFP,attP}, pBac{hr5iei::ZsGreen-p10}*, and *pBac{ie1::mCherry, ie-UAS-Gal4Δ::mCD8-GFP}* were respectively gifts from David Stern (Addgene plasmid # 86860 ; RRID:Addgene_86860), Tim Harvey-Samuel, and David Miller. Bacterial cells containing plasmids were grown in 50 mL liquid cultures of Luria-Bertani broth with 100 µg/mL ampicillin, and plasmid DNA was subsequently purified with the ZymoPURE II Plasmid Midiprep Kit (Zymo Research, #D4200). Plasmids were eluted in 120 µL of elution buffer, yielding concentrations of 620 - 1190 ng/µL of plasmid DNA.

*Hyperactive Piggybac* transposase mRNA (*HyPBase*) was prepared in vitro from the *pGEM-T_hyPB^apis^* plasmid, a kind gift from Martin Beye (Otte et al., 2018). This plasmid was modified using site-directed mutagenesis to render it compatible with 5’ capping of the mRNA with CleanCap Reagent AG (New England Biolabs). To do this, we generated a G→A single nucleotide mutation immediately 3’ of the T7 promoter TATA sequence using the Q5 Site-Directed Mutagenesis Kit (New England Biolabs, #E0554S) following the manufacturer’s recommendation, with the exception that the KLD reaction incubation time was increased from 5 min to 60 min. The primers for this mutagenesis (*hypB_AG_Q5F:* 5’-ACTCACTATAaGGCGAATTGG-3’, and *hypB_AG_Q5R:* 5’-CGTATTACAATTCACTGGC-3’) were designed using the NEBaseChanger online tool. The resulting plasmid was introduced into DH5a *E. coli* competent cells (Zymo Research, #T3007), amplified in 10 mL liquid culture, and purified using the QIAprep Spin Miniprep Kit (Qiagen). The mutation was validated by whole plasmid sequencing through Primordium Labs.

This modified plasmid was then linearized with NcoI-HF, and purified using the DNA Clean & Concentrator-25 kit (Zymo Research). Around 500 ng of linearized template was used for transcription, poly-adenylation and capping using the HiScribe T7 mRNA Kit with CleanCap Reagent AG (New England Biolabs, #E2080S), and mRNA was purified using the MEGAClear Transcription Clean-Up Kit (Thermo Fisher Scientific). The purified mRNA was eluted in 50 µL of pre-heated elution buffer provided with the kit, and incubated on the filter column for 10 min in a 65°C oven. The eluted RNA was re-applied to the column and the step was repeated. After quantification with Nanodrop (Thermo Fisher Scientific), the transposase mRNA was aliquoted into one time use batches of >1000 ng/µL to avoid freeze-thawing and stored at −70°C. Injection mixes were freshly prepared before injection and consisted of 400 ng/mL transposase mRNA, 200 ng/mL donor plasmid, and 0.05% Phenol Red (1:10 dilution of a 0.5% cell-culture grade Phenol Red solution; Sigma-Aldrich, St. Louis, MO, USA), brought to a 5 µL final volume with 1x Bombyx injection buffer (pH = 7.2-7.4, 75 µM NaH₂PO₄, 425 µM Na₂HPO₄, 5 mM KCl).

### Embryo microinjections

Microinjections of embryos at the syncytial stage followed published protocols (Heryanto, Mazo-Vargas, et al., 2022; Martin et al., 2020) with species-specific adjustments. Microinjection needles were prepared using a PC-10 gravity puller (Narishige International) on heat settings 58-62, using borosilicate capillaries (#18100F-3, World Precision Instruments). Syncytial embryos were injected within 15–120 min after egg laying (AEL): 15–35 min for *P. interpunctella,* and 20-120 min for *J. coenia*, *V. cardui*, *A. incarnata*, and *A. gemmatalis.* To prevent desiccation, *P. interpunctella* embryos were sealed with cyanoacrylate glue (GH1200 Super Glue All Purpose with Anticlog Cap) at the injection site, incubated at 28°C for 24 hr under high humidity conditions in a closed tupperware containing a wet paper towel, and then moved to a rearing container with diet at 28°C. Similarly, microinjected embryos of *J. coenia*, *A. incarnata*, and *A. gemmatalis* were incubated in humid chambers at 28°C for 24 hr, or at 28°C for *V. cardui*, but do not require manual sealing with glue. Survival and phenotypes were scored at hatchling, larval, pupal, and adult stage (Supporting Information Table S2).

### Generation of CRISPR germline mutant lines

Stable, germline-carrying lines of *scarlet* knockout mutants were established in *V. cardui, J. coenia*, *P. interpunctella*, and *A. gemmatalis* by in-crossing multiple G_0_ crispants with mosaic eye coloration defects. In all four cases, this random in-crossing scheme was sufficient to generate a large proportion of homozygous-recessive mutants at the G_1_ generation, which were then selected for further in-crossing and establishment of stable lines lacking WT alleles. Similar strategies were used to attempt the generation of stable *white* KO lines in *V. cardui*, *J. coenia*, and *A. gemmatalis*.

### Brightfield macrophotography

Images of wild-type and crispant individuals were captured using a Keyence VHX-5000 and Keyence VHX-7000 digital microscope, or a Nikon D5300 Nikon camera mounted on a stereomicroscope. Notably, *A. gemmatalis* moths exhibit light-adaptive changes in their eyes, where pigment granules shift position to manage light sensitivity (Nordström & Warrant, 2000), causing color shift from light grey to dark brown. For each adult eye comparison we selected the wild-type individuals that were under the same light conditions as the ABCG transporter mutants.

### Fluorescent screening and imaging

Fluorescent screening of eggs, larvae, and pupae was performed using an Olympus SZX16 stereomicroscope equipped with a Lumencor SOLA Light Engine SM 5-LCR-VA light source, and imaged via an Olympus DP73 digital color camera mounted on a trinocular tube. Chroma Technology filter sets were used for the detection and separation of fluorescent markers as follows: EGFP (ET470/40x, ET510/20m) for screening EGFP and ZsGreen, EYFP (ET500/20x, 535/30m) for screening EYFP as well as EGFP and ZsGreen with reduced autofluorescent background, and AT-TRITC-REDSHFT (540/25x, 620/60m) for detecting mCherry. For pupal screening, individual fifth instar larvae were separated and allowed to pupate in small plastic cups. Each pupa was then examined for a positive fluorescent signal, either in the developing eye when examining *pBac{3xP3::EYFP}* or in the entire body when screening for *ie1*-driven activity with *pBac{hr5iei::ZsGreen-p10}* and *pBac{ie1::mCherry, ie-UAS-Gal4Δ::mCD8-GFP}*.

### Genotyping

For verification of DNA lesions, genomic DNA was extracted from the thorax of individuals using the Quick-DNA Tissue/Insect Miniprep Kit (Zymo Research, #D6016). Fragments of 700-900 bp were amplified by PCR using primers flanking the guide RNA target sites, and an M13 universal sequence added to the forward primers (**Supporting Information Table S1),** and purified using the Zymo DNA Clean & Concentrator-25 kit eluted with 15 μL of DNA elution buffer. Long-read sequencing was performed at Azenta, and analyzed with the WF-AMPLICON workflow of the EPI2ME software (**Supporting Information Fig. S8**).

## RESULTS

### Pleiotropic effects of *white* knockouts on pigmentation and survival

To investigate how ABC-mediated trafficking influences pigmentation in eyes and integument, we first mutated the *white* gene in the nymphalid species *V. cardui, J. coenia,* and *A. incarnata,* and in the moth *A. gemmatalis* using CRISPR-Cas9. We injected and monitored the pigmentation of G_0_ individuals throughout development, from early larval stages through pupation, and compared their coloration to similarly staged wild-type controls. Across the four species, our mutagenesis efforts successfully generated G_0_ mosaic individuals with a range of visible larval pigmentation and integumentary phenotypes at rates of 16%, 24%, 45% and 50% in *V. cardui*, *J. coenia*, *A. incarnata*, and *A. gemmatalis* respectively (**Figure 1; Supporting Information Fig. S1, Table S2**). The G_0_ *white* mosaic mutants displayed a range of phenotypes, including patches of pigmentation loss or lighter pigmentation in the larval and pupal stages (**Figure 1A, B, D, E, G, H, J; Supporting Information Fig. S1B**). The three butterfly species also presented misshaped larval bristles and changes in the coloration of the pupal epidermis. *V. cardui* and *J. coenia* (not shown) developed exaggerated bulges at the posterior end of the larvae, typically on the dorsal side of abdominal segment 8 (A8), that were more pronounced than in wild type (**Supporting Information Fig. S1B**). In *A. incarnata*, we further observed defects in egg coloration, where the characteristic brownish–red pigmentation that develops after 24 hours failed to appear in the mutant clones (**Supporting Information Fig. S2**). Most notably, butterfly larvae with extensive mutant regions appeared sickly throughout development, with a seemingly darker exterior outside the lighter mosaic patches (**Supporting Information Fig. S1B**). This phenotype appears to be due to molting defects during transitions between larval instars and the inability to fully shed the old larval integuments. Finally, *white* mosaic knockouts resulted in adult eye tissues lacking any pigmentation (**Figure 1C, F, I, L**), as typically observed in other insects. Adults did not show pigmentation defects in wing scales or in other regions than their eyes.

**Figure 1.**
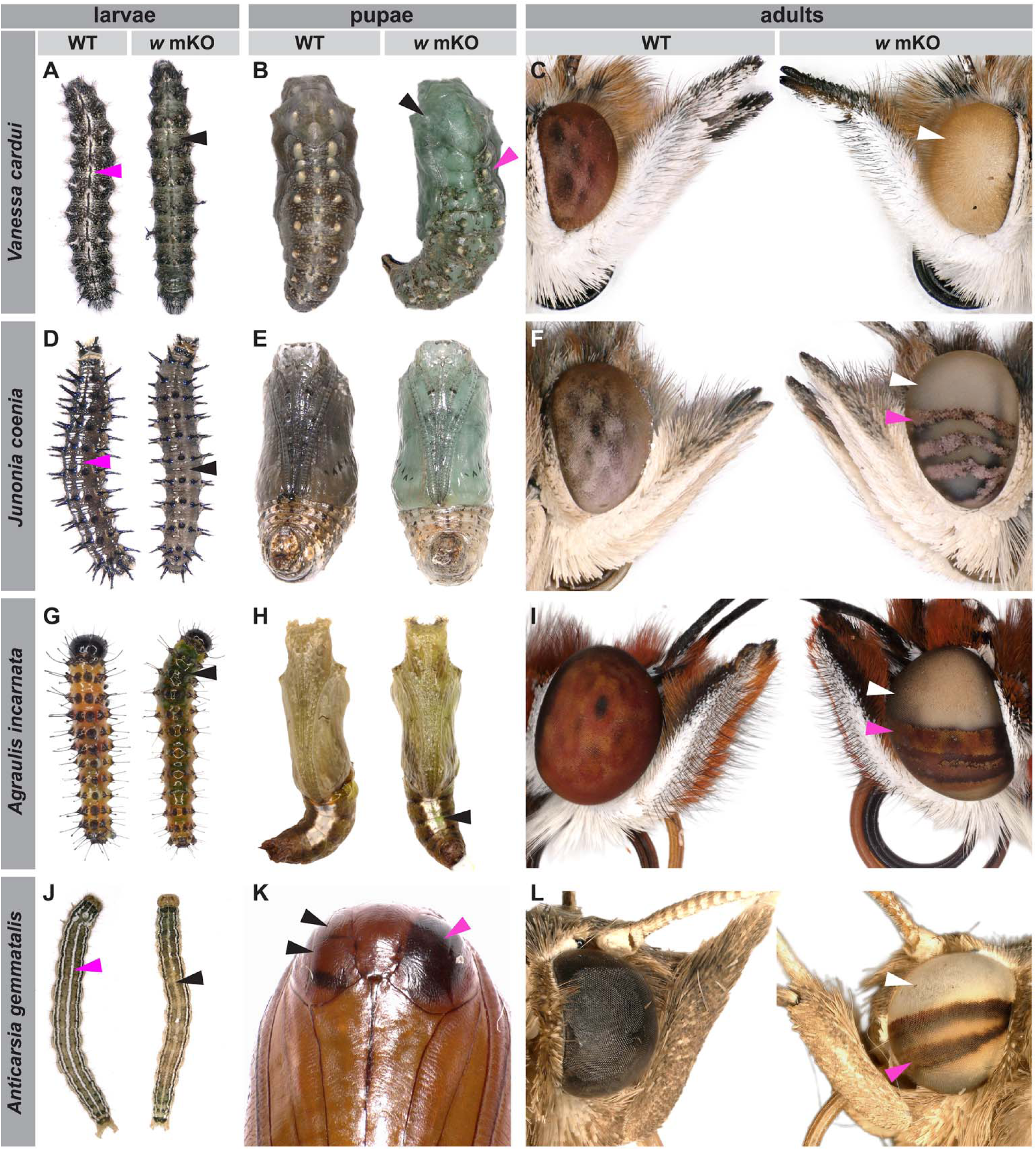
CRISPR-Cas9 disruption of the *white* gene alters pigmentation during larval, pupal, and adult stages in Lepidoptera. For each species, the larva, pupa, and adult of both wild–type (WT, left) and mosaic knockout (mKO, right) individuals are shown. Magenta arrows indicate characteristic WT pigmentation, whereas white and black arrows highlight mosaic clones resulting from *white* gene disruption. **(A-C)** In *V. cardui*: (**A**) *white* mKO L4 larva has lost the white and dark grey pigmentation in mutant clones visible down the dorsal midline (black arrowhead). (**B**) Dorsal view of a *white* mKO pupa at 0-4 hr APF (After Puparium Formation) of pupation compared to WT shows disruption of the dark green/brown pigmentation as well as the white to yellow coloration of the protrusions. (**C**) Adult mKO eyes are discolored compared to the reddish-brown wild-type eyes. (**D-E**) In *J. coenia:* (**D**) large mutant clones in a *white* mKO L3 caterpillar show loss of overall dark coloration and white patterns along the midline. (**E**) Ventral view of a *white* mKO pupa at 0-4 hr APF displays lighter coloration compared to the dark green WT (**F**) depigmented mosaic clones in adult eyes compared to the brown-eyed wild type. (**G-I**) In *A. incarnata*: (**G**) *white* mKO larva appears a brighter green than WT, with plant tissue visible in the gut, due to increased tissue transparency (**H**) mKO pupa at 0-4 hr APF is a brighter green compared to the WT due to tissue transparency (**I**) adult mKO eyes are discolored compared to the reddish-brown wild-type eyes. (**J-L**) In *A. gemmatalis:* (**J**) *white* mKO larvae and (**K**) pupa with mosaic clones (magenta arrowhead: dark, wild-type pigment, black arrowheads: depigmented, mutant clones). (**L**) Mutant clones in adult mKO eyes appeared white (white arrowhead) compared to the brown wild type (magenta arrowhead).

In previous work, we determined that CRISPR G_0_ somatic mutants often carry mutations in their germlines, and crossing them can allow the generation of compound heterozygous mutants in the G_1_ generation (Fandino et al., 2024; Livraghi et al., 2024; Mazo-Vargas et al., 2022). In an attempt to generate *white* homozygous-mutant lines in *V. cardui, J. coenia*, and *A. gemmatalis*, we performed random matings of G_0_ crispant individuals. These germline transmission attempts, including three replicated trials in *V. cardui*, all produced fertile offspring, but failed to generate pigmentation phenotypes in G_1_ late larvae, pupae, or adults, indicative of deleterious pleiotropy of the *white* mutations in these species. Of note, while we recovered G_1_ offspring with mutant phenotypes in early stage larvae of *J. coenia* and *V. cardui,* these individuals did not survive past the second instar. In *V. cardui*, for example, ten G_1_ larvae displayed clear mutant phenotypes at the first instar (**Supporting Information Fig. S1C**), but all failed to reach pupation and died during early larval development. These individuals exhibited pronounced molting defects including a double integument appearance, with new bristles visible beneath the unshed integument, and similar to previously described *white* mutants in *Helicoverpa* caterpillars (Khan et al., 2017), the retention of the previous head capsule following a larval molt (**Supporting Information Fig. S1, D-H**). In the case of *A. gemmatalis,* random G_0_ crossing produced offspring without visible phenotypes. In summary, *white* mosaic knockouts reliably yielded pigmentation phenotypes in the four lepidopteran species we tested, but these mutations failed to produce viable mutant offspring in three of the species. In particular, the lethal effects of *white* germline mutations, manifesting as molting defects in nymphalid butterflies, highlight that *white* may have pleiotropic roles beyond eye and integument pigmentation. Because White functions as an obligate heterodimer with distinct ABC transporters that mediate uric acid and ommochrome transport, we next asked whether disrupting these partner proteins might separate pigmentation phenotypes from the severe developmental defects observed in *white* mutants. Next, we targeted the *ok* gene, which encodes a proposed partner of White (Fujii & Banno, 2019; J. Lee et al., 2018; Ninomiya et al., 2006).

### Conserved role of *ok* in purine-derivative trafficking in Lepidoptera

We disrupted the ABCG transporter gene *ok* in *V. cardui*, *A. incarnata*, and *A. gemmatalis* to test its role in integument and eye pigmentation across life stages, recovering G₀ mosaic individuals at 50%, 56%, and 25%, respectively. In all three species, *ok* mosaic knockouts consistently reduced or eliminated the opaque white pigmentation of larval and pupal integuments, producing translucent or depigmented effects (**Figure 2**). A darker tint was consistently observed in pupae of *V. cardui* and *A. incarnata* following *ok* disruption, in contrast to the characteristic green/blue cast seen after *white* gene perturbation (**Figure 2B, E)**. Based on parallel evidence in silkmoths (Tamura & Akai, 1990; Wang et al., 2013), these *ok* mutant phenotypes are attributable to a loss of uric acid, a white opaque pigment and excretory product that is commonly stored in the epithelial cells of butterfly and moth caterpillars (Fujii & Banno, 2019; J. Lee et al., 2018; Lhonoré et al., 1980; Ninomiya et al., 2006)

**Figure 2.**
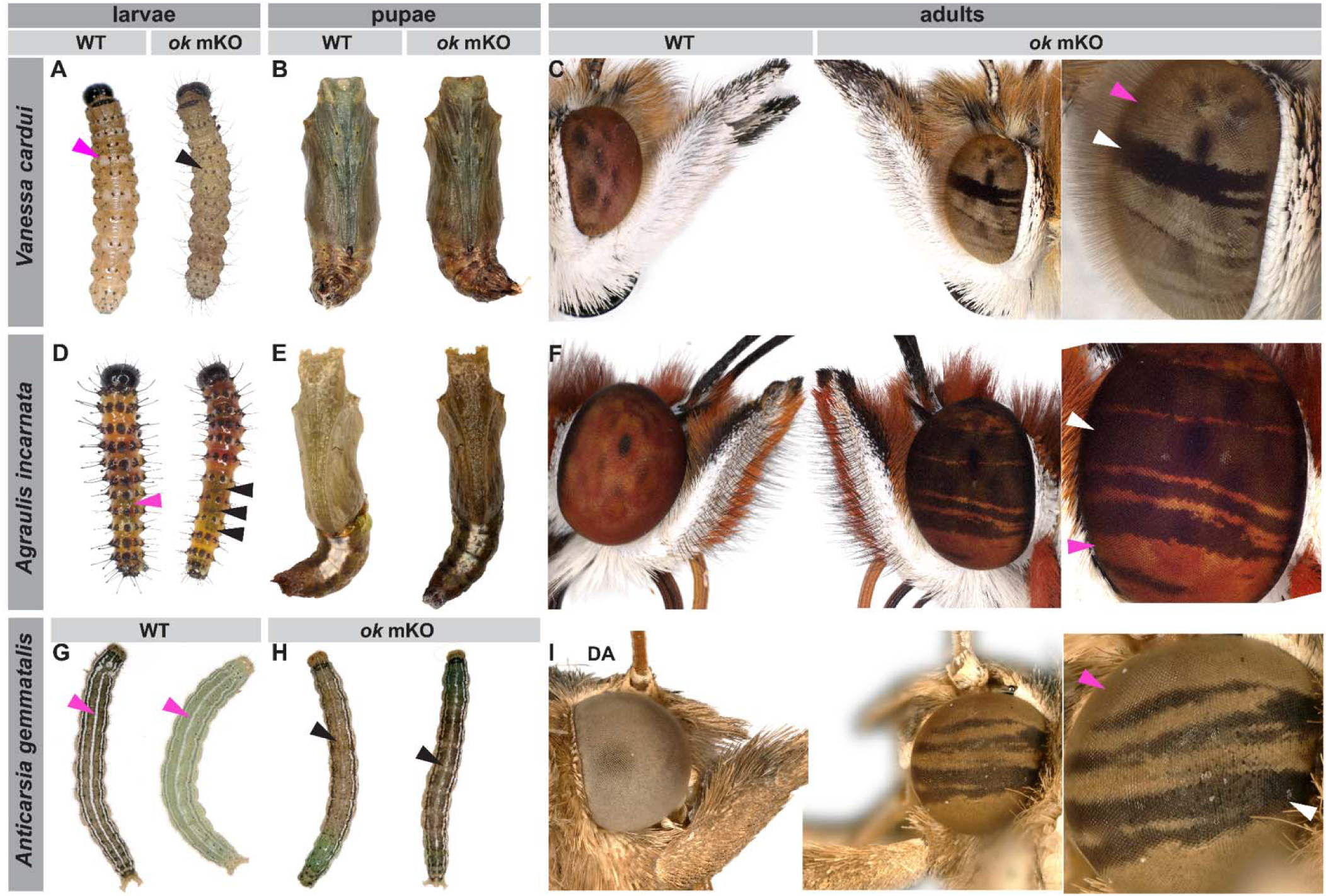
CRISPR-Cas9 disruption of the *ok* gene alters pigmentation during larval, pupal, and adult stages in Lepidoptera species. For each species, the larva, pupa, and adult of both wild–type (WT, left) and mosaic knockout (mKO, right) are shown. Magenta arrowheads indicate characteristic WT pigmentation, and white or black arrows highlight mosaic clones resulting from *ok* gene disruption. (**A-C**) *V. cardui*: (**A**) *ok* mKO larva has lacking white patches (black arrow) (**B)** *ok* mKO pupa at 0-4 hr APF has an overall darker appearance compared to WT (**C**) *ok* mutant clones in adult eyes are darker than the WT coloration (**D-F**) *A. incarnata*: (**D**) *ok* mKO larva missing white spots in the posterior segments (**E)** *ok* mKO pupa at 0-4 hr APF appears overall darker than stage matched WT (**F**) *ok* mutant clones in adult eyes are dark brown compared to the orange-red WT eyes (**G-I)** *A. gemmatalis:* (**G)** WT larvae, dark (left) and light (right) morphs. **(H)** The white stripe down the midline is disrupted in *ok* mKO larvae. **(I)** *ok* mKO adult eye clones appear darker than the dark adapted WT eye (DA : dark-adapted, *i.e.* with outer brown pigments retracted and procuring a light coloration to the outer surface of the eye).

In nymphalid butterflies, individuals with extensive *ok* mutant mosaics frequently displayed compromised larval survival, including slowed growth and molting difficulties (**Supporting Information Fig. S3**). Although we did not establish stable *ok* lines in the nymphalids, these observations echo the non–pigmentary integument defects we documented for *white* loss-of-function and suggest that Ok, like White, contributes to molting physiology in at least some Lepidoptera. Together, these outcomes indicate that while uric acid consistently contributes to white pigmentation of larval and pupa integuments and opacity, the magnitude and visibility of its loss vary by lineage due to differences in baseline urate allocation and co–occurring pigment systems.

As previously observed in *Helicoverpa* moths (Khan et al., 2017), the adults of all three species exhibited darker eye pigmentation in *ok* mutant clones compared to wild-type eye tissues (**Figure 2C, F, I**). These dark mutant clones suggest that *ok* is required for the deposition of light-colored screening pigments in lepidopteran eyes, possibly of the pteridine class based on a previous report (Khan et al., 2017). In *A. gemmatalis*, the ocular effects of *ok* mKOs were visible when the crispants were exposed to low-light conditions before imaging. In nocturnal moths, low-light conditions trigger a withdrawal of dark screening pigments from the light path that maximize the sensitivity of the retina (Berry, 2022; Dreisig, 1981). When dark-adapted, WT eyes take a light-grey colorations, and crispants for *ok* showed darkened clones. In this model, pterins act as eye pigments and their removal in *ok* crispants unmasks the presence of ommochromes, resulting in a darkening effect. The identification of these pigments, and their histological localization within the complex anatomy of the lepidopteran compound eye, will require further investigation. Similar to *white* mutants, we did not observe any scale pigmentation phenotypes in the *ok* mosaic knockouts.

### Conserved roles of *scarlet* in eye and integumentary ommochrome pigmentation

We next disrupted the ABCG transporter gene *scarlet* in five lepidopteran species, *J. coenia, V. cardui, A. incarnata, P. interpunctella,* and *A. gemmatalis,* to test whether impairing ommochrome transport alters pigmentation across life stages without causing the lethality observed in *white* knockouts. Across the five species, we generated G_0_ mosaic individuals with a range of pigmentation and integumentary phenotypes. Unlike *white*, disruption of *scarlet* did not cause any deleterious effects, and we obtained stable lines of *st* knockouts in *V. cardui, J. coenia*, *A. gemmatalis,* and *P. interpunctella*.

In the three nymphalid butterflies, knockout of *scarlet* resulted in visibly lighter larval and pupal coloration (**Figure 3**) especially in first to third instars. Pupae at 0 hr APF displayed a light blue-green color hemolymph (**Figure 3B, E, H**), and lighter integuments at all other pupal stages. All five species exhibited strikingly light eye phenotypes compared to their dark wild-type eyes (**Figure 3**), which was also apparent in the developing pupal eyes of the moths *A. gemmatalis* and *P. interpunctella* (**Figure 3K, N).** In nymphalid butterflies, *st* knockouts exhibited a light beige adult eye coloration, consistent with the loss of ommochrome pigments. *A. gemmatalis* and *P. interpunctella st* knockout adults, however, displayed yellow to rosy and golden colored eyes, respectively, indicating the presence of pteridine pigments (**Figure 3L, O**).

**Figure 3.**
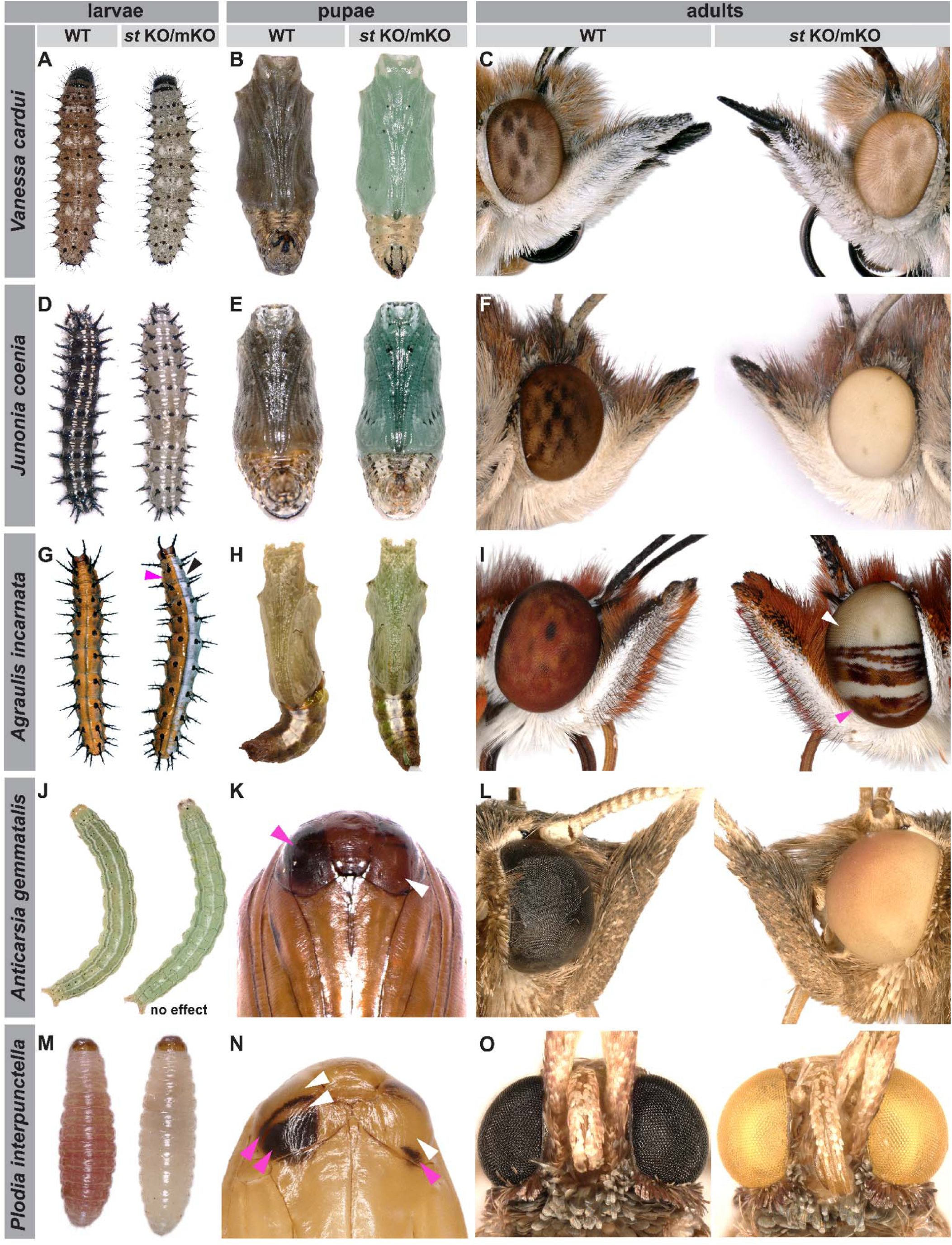
CRISPR-Cas9 disruption of the *scarlet* gene alters pigmentation during larval, pupal, and adult stages in Lepidoptera. For each species, we show the larva, pupa, and adult of both wild–type (WT, left) and knockout or mosaic knockout (KO/mKO, right) individuals. Magenta arrows indicate characteristic WT pigmentation, while white and black arrows highlight mosaic clones resulting from *st* gene disruption. Pupae in B,E,H are 0-4 hr APF. **(A-C)** stable G3 *st* KO line in *V. cardui*. (A) L2 *st* KO caterpillars (right) display an overall lighter body coloration compared to brown-pink WT *V. cardui* larvae at this stage (left). (B) *st* KO pupa display a light greenish-blue hemolymph (right) compared to the dark olive WT (left). (C) Adult st KO eye (white arrowhead, right) exhibits a light beige coloration, much lighter than the brown pigmented WT eyes (magenta arrowhead, left). These phenotypes are similarly observed in the **(D-F)** stable *st* KO line in *J. coenia.* **(G-I)** *A. incarnata ok* mKO **(G)** bilateral mutant larva shows loss of orange coloration (black arrow). (H) *st* mKO pupa displays a brighter green compared to WT (I) mosaic *st* clones in adult eyes appear depigmented **(J-L)** *A. gemmatalis:* (**J**) no effect of *st* KO is observed in larval coloration (stable line, G_3_) **(K)** G_0_ pupa with lighter colored mosaic eye clones (white arrowheads) **(L)** *st* KO G_3_ adult eye with a yellow to rosy color, instead of the dark brown WT phenotype. **(M-O)** *P. interpunctella:* (**J)** *st* KO larva with depigmented integument **(N)** G_0_ pupa showing lighter colored mosaic eye clones (**O**) *st* KO adult eye with a golden coloration compared to the dark brown WT eye.

Knockouts of *white* in *Plodia* are viable and have been used as a genetic background for generating transgenic lines (Heryanto, Mazo-Vargas, et al., 2022; Shodja et al., 2025). However, disruption of *white* eliminates pigmentation throughout the larval body, including the otherwise dark brown testes, thereby preventing reliable sex determination at larval stages. In *scarlet* mutants, while the larval integument exhibits a transparent appearance, the testes change from the wild-type dark brown to light yellow, allowing to distinguish between males and females (**Supporting Information Fig. S4**). Notably, all *scarlet* mutants had phenotypically wild–type scales, making the *scarlet* mutation an optimal genetic background for transgenesis when investigating wing–pattern or scale–specific phenotypes of interest.

### The bright orange of *A. incarnata* larvae emerges from a combination of ommochrome and uric acid

The mutant larvae of *A. incarnata* exhibited striking integumentary effects and shed new light on the molecular basis of caterpillar coloration. wild-type *A. incarnata* caterpillars display a bright orange coloration, with longitudinal stripes positioned laterally and on the dorsal midline that appear darker (**Figure 4A, C, C’**). Strikingly, mosaic knockouts of *scarlet* resulted in transformation of orange to white integument. Upon closer examination of these whitened tissues, we noticed a heterogeneous distribution of an opaque white substance in the areas normally colored bright orange (**Figure 4B, E**). In addition, areas that are normally darker, such as the lateral lines or the proximal sections of the legs and prolegs, lost their brown tint in *scarlet* mutants, and appeared more transparent (**Figure 4B, D, E**). Thus, the *A. incarnata* larval integument contains a ubiquitous ommochrome pigment, which when combined with a white opaque pigment, produces a vivid orange effect. Mutants for *ok* suggest that this white pigmentation is likely uric acid. Indeed, *ok* mutant clones appeared darker and browner, consistent with the loss of the white pigment while retaining the Scarlet-dependent ommochrome tint (**Figure 4F, F’**). Finally, *white* mosaic KOs rendered the caterpillar integument completely transparent, revealing the tracheal system and the developing wing disc through the clear thoracic integument (**Figure 4G, G’**), or otherwise allowing the coloration of the gut contents, fat bodies and hemolymph to show through (**Supporting Information <u>Video S1</u>**). Together, these data lead to a model where White-Scarlet dimers are required for ommochrome pigmentation across the larval integument, while White-Ok dimers are required for accumulation of uric acid in large patches of the caterpillar (**Figure 4H**). The combination of ommochrome and uric acid results in bright orange coloration.

**Figure 4.**
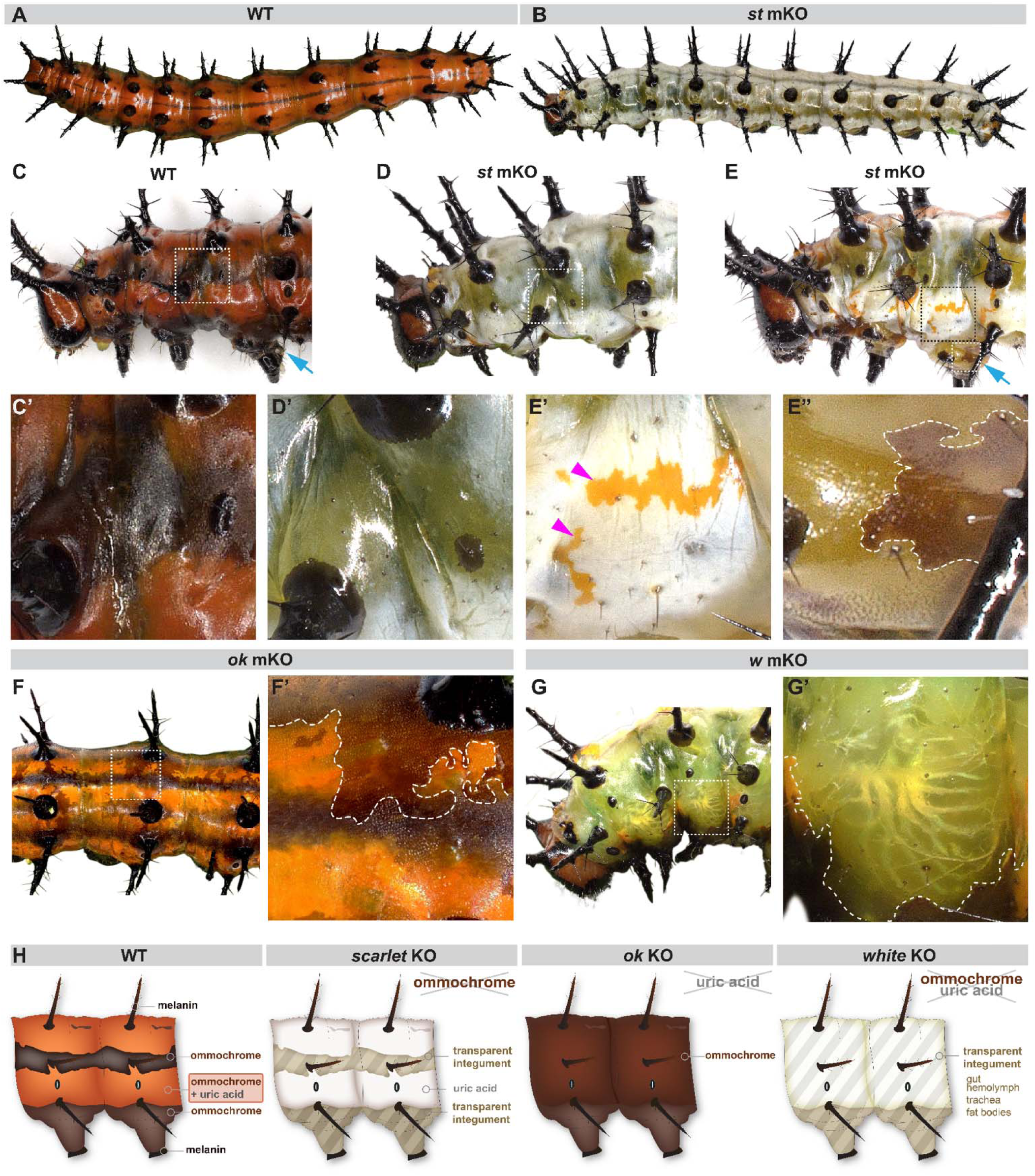
Effects of *white*, *scarlet,* and *ok* knockouts on larval integument coloration in *A. incarnata.* (**A**) Dorsal view of wild type (WT) *A. incarnata* larva with bright orange coloration (**B**) Lateral view of a *scarlet* mKO larva with an overall white appearance. (**C-E**) Lateral views of anterior segments: (**C**) WT larva with bright orange coloration, dark longitudinal stripes, and dark legs (blue arrow). The dark longitudinal stripe and legs became transparent in *scarlet* mKO individuals (**D, E,** compare **D’ to C’,** and arrows in **C** and **E, E”**). The orange coloration appears white in *scarlet* mKO **(D,E,E’)** . (**F**) *ok* mKO clones are darker than the WT orange. (**G**) *white* mKO larva has large transparent clones in regions that are orange or dark in WT individuals. Trachea and developing wing disc (**G’**) are visible through the thoracic integument. (**H**) Summary of knockout effects and inferred contributions of different pigment types.

A potential complication to this model is that *scarlet* mosaic mutants, in some occurrences, produced a small proportion of patches with translucent integument, resembling the phenotypes observed in *white* knockouts (**Supporting Information <u>Video S2</u>**). We propose that in these clones, *scarlet* alleles generated by CRISPR mutagenesis produced truncated Scarlet proteins that interfered with White function, potentially resulting in partial disruption of urate granule formation. Because a single CRISPR target site was used to disrupt *scarle*t, about a third of indel mutations (multiple of 3 bp in size) are expected to preserve the reading frame while altering protein structure, which could underlie these mosaic phenotypes via a dominant-negative effect.

### Comparative analysis of co-transporter KOs in larval integuments

We next examined how combinations of pigment classes contribute to larval pigmentation in three additional species. *V. cardui* caterpillars have a characteristic reddish-brown coloration at the L2-L3 (second and third larval instar) stages, with two white stripes along the midline and white spots on either side of the midline in every other segment (**Figure 5A**). In *white* mKOs, both the reddish-brown pigmentation and white stripes and spots were disrupted (**Figure 5B**), resulting in either transparent or de-pigmented mutant clones. Mosaic perturbations of *ok* yielded a marked reduction of the white spots and stripes on the larval integument (**Figure 5C**), consistent with a deficiency in uric acid. Last, stable germline mutants for *scarlet* (homozygous mutants at the G_1_ and G_2_ generations) showed a complete loss of reddish hues, seen as white-colored larvae, but left white patterns unaffected (**Figure 5D**). In a nutshell, phenotypes were consistent with a loss of uric acid in *ok*, a loss of ommochromes in *scarlet* mutants, and a loss of both classes of pigments in *white* mutants.

**Figure 5.**
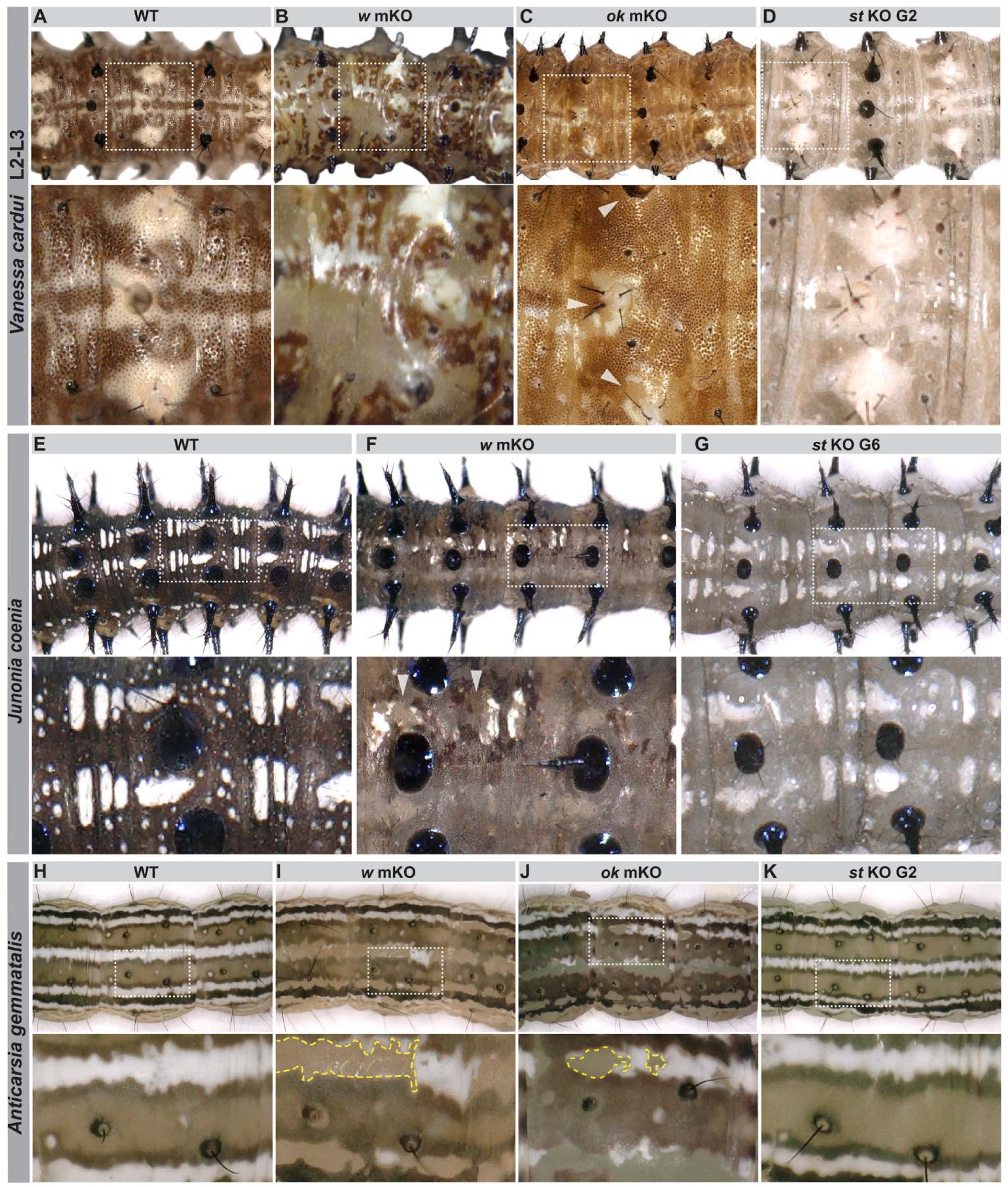
Effects of *white*, *ok*, and *scarlet* knockouts on larval integument coloration in *V. cardui*, *J. coenia*, and *A. gemmatalis*. Larvae are oriented with the anterior to the left in all panels. (A-D) *V. cardui* second to third instar larvae (**A**) Dorsal view of a WT caterpillar with reddish-brown coloration (shade of brown varies among WT individuals), two parallel white stripes along the midline and two white spots on both sides (inset). (**B**) Both brown and white pigmentation are disrupted in a *white* mKO individual, generating transparent clones. (**C**) Disruption of *ok* impaired white pigment deposition (arrowheads in inset), while (**D**) in stable *scarlet* mutants only the ommochrome based reddish-brown coloration was affected, resulting in a lighter colored larva with intact white pigments. (E-G) *J. coenia* third instar larvae: (**E**) WT larva has a dark grey-brown color with white patterns along the dorsal midline. (**F**) *white* mKOlarvae show lighter colored clones with impaired brown and white pigment deposition (arrowheads mark example clones). (**G**) Stable *scarlet* KO displays an overall lighter coloration without affecting white patterns. **(H-K)** Dorsal views of *A. gemmatalis* larvae: (**H**) representative WT larvae with green pigmentation and white stripes. Both (**I**) *white* mKO, and (**J**) *ok* mKO disrupted the white stripes but not the green pigmentation. (**K**) Neither green pigmentation nor white striping was affected in stable *scarlet* KO individuals (G_2_).

In *J. coenia* larvae at the L3 stage, mosaic KOs of *white* were similar to *V. cardui* as they resulted in mutant clones with a reduced opacity, lighter pigmentation, and loss of white patterns (**Figure 5E, F**). While we did not test for the function of *ok* in this species, it can be inferred that these white patterns also require a functional White-Ok heterodimer. Here as well, stable *scarlet* mutant lines had an overall lighter phenotype, indicative of ommochrome deficiency, and maintained their white patterns along the midline (**Figure 5G**).

*A. gemmatalis* larvae display significant variations in integument coloration, spanning light green, to brown, to nearly black, in part due to a plastic response to larval crowding conditions (Silva et al., 2013). White stripes that run laterally and on the dorsal midline are invariable among these morphs, and were the only pattern elements disrupted upon mosaic knockouts of both *white* and *ok* (**Figure 5H-J**). No effects were detected in *scarlet* mutants, including in homozygous-mutant individuals at the G_2_ generation (**Figure 5K**). These data suggest that *A. gemmatalis* use uric acid in their white elements, but in contrast with nymphalids and *Helicoverpa* moths (Khan et al., 2017), other aspects of their coloration are not dependent on ommochromes.

### White-Ok and White-Scarlet pairs contribute to pupal integument opacity and coloration in nymphalids

To assess the effects of ABC transporter knockouts on pupal integuments, we compared all the integuments at 0-4 hr APF (after pupal formation) prior to the complete cuticle sclerotization. The cuticle of wild-type pupae of *V. cardui* rapidly mature into an opaque beige coloration, and are adorned with orange-gold protrusions on the dorsal side (**Figure 6A**). Mosaic knockouts of *white* severely disrupted these features, resulting in patches of transparent integument dominated by the blue-green hue of the underlying hemolymph pigments, and losing the orange-gold coloration of their dorsal protrusions (**Figure 6B**). Mutation of *ok* resulted in mosaic darkening of the integument due to loss of white pigmentation, likely from uric acid deficiency (**Figure 6C**). In contrast, *scarlet* mutation generated overall lighter color pupae with a blue-green tint (**Figure 6D**), consistent with the loss of a ubiquitous ommochrome pigment that would normally mask the pigmentation of the pupal hemolymph. Orange-gold protrusions appeared a paler color in *scarlet* knockouts, suggesting this feature is a combination of ommochrome, uric acid, and an additional pigment, such as pterins. Similar results were obtained in the larvae of *A. incarnata* for *white*, *ok*, and *scarlet* (**Figure 6E-H**), and in the pupae of *J. coenia* for *white* and *scarlet* (**Supporting Information Fig. S5**). We did not detect defective integumentary phenotypes in *white* and *scarlet* loss-of-function in the pupae of *A. gemmatalis* and *P. interpunctella*, suggesting their dark brown coloration is independent from purine or ommochrome derivatives. Taken together, these results show that the pupae of nymphalid butterflies include an ommochrome-based brown component, as well as a white component (likely uric acid) that contributes to their opacity.

**Figure 6.**
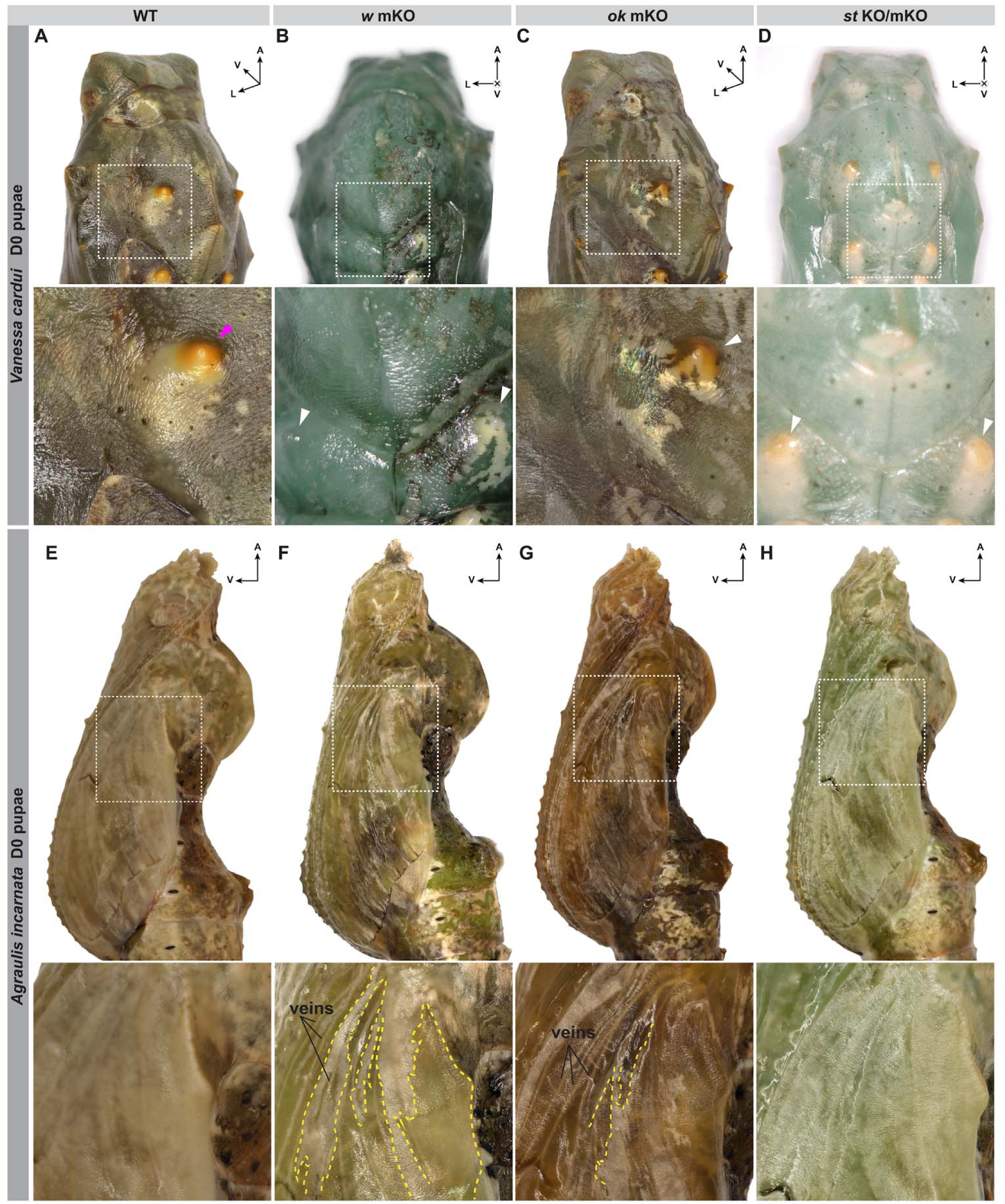
Effects of *white*, *ok,* and *scarlet* knockouts on pupal integument coloration and opacity in nymphalids. **(A-C)** *V. cardui* pupae at 0-4 hr APF: (**A**) Dorsolateral view of a WT pupa with olive green coloration and white patches. Inset shows orange-gold protrusion (magenta arrowhead). (**B**) Dorsal view of *white* mKO pupa showing transparency of the left half, including a dorsal protrusion, and loss of orange-gold coloration in the right protrusion, with white-colored clones consistent with uric acid deposition and absence of ommochrome (white arrowhead). (**C**) Dorsolateral view of an *ok* mKO individual with defects in white patches. Inset shows loss of uric acid based white coloration in a dorsal protrusion and the presence of dark mosaic clones. (**D**) Dorsal view of a stable *scarlet* KO individual shows overall lighter coloration, and gold protrusions appear paler. **(E-H)** *A. incarnata* at 0-4 hr APF, all lateral views: (**E**) Brown WT pupa with white-colored patches creating opacity. Opacity is disrupted in (**F)** *white* mKO and (**G**) *ok* mKO, but not in (**H**) *scarlet* mKO; *white* mKO and *scarlet* mKO individuals appear overall lighter in color.

### The *scarlet* mutant background enables detection of fluorescent proteins for transgenesis

Having successfully generated *scarlet* mutant lines in both *J. coenia* and *V. cardui,* we next evaluated their fluorescence screenability for transgenesis. Using *piggyBac* mediated transposon transgenesis (Heryanto, Mazo-Vargas, et al., 2022), we tested three donor plasmids to assess different drivers and fluorescent proteins. Namely, we tested (1) an eye and glia specific promoter (*3xP3*) driving enhanced yellow fluorescent protein (EYFP) (hereafter *pBac[3xP3::EYFP, attP]*), and (2) two separate plasmids with a baculovirus immediate-early (ie) promoter expected to have broad, non-tissue specific activity (Guarino & Dong, 1991; Masumoto et al., 2012), driving either green (*pB[hr5/ie1::ZsGreen])*, or and red (pB[hr5/ie1::mCherry]) fluorescent proteins. For both species, we microinjected eggs between 1-2 h AEL across multiple injection trials. Using the *pBac[3xP3::EYFP, attP]* plasmid, we detected fluorescence in the developing G0 larval and pupal ocelli and eyes (**Supporting Information Fig. S6, Table S3**) When using the *pB[hr5/ie1::ZsGreen]* and *pB[hr5/ie1::mCherry]* plasmids, we detected localized patches of fluorescence in the G_0_ developing pupae, in a spatially random patterns (**Supporting Information Fig. S7**). Notably, detection of fluorescence was markedly easier in the *scarlet* mutant background; fluorescent patches in each individual were more abundant, spatially larger, and qualitatively brighter compared with those observed in a wild-type background.

Overall, *scarlet* mutant lines enabled screening of fluorescence through the developing tissues in G_0_ individuals, providing a practical approach for identifying transgenic events. In efforts to establish stable transgenic lines, we implemented various G_0_ crossing strategies; however, none yielded transgenic offspring.

## DISCUSSION

### Conserved role of White-Scarlet in eye ommochrome pigmentation across Pancrustacea

We showed that both *white* and *scarlet* knockouts result in eye coloration phenotypes. A review of the literature reveals that these two genes provide robust eye marker effects on a broad set of insect species where these genes have been knocked-out or knocked-down (**Figure 7A, B**). Based on the *Drosophila* classic model that White-Scarlet heterodimers transport an ommochrome pigment precursor (Sullivan et al., 1979; Tearle et al., 1989), *scarlet*-deficient insects can show residual pigmentation that indicates the presence of other eye pigments, as seen in Lepidoptera, Coleoptera, and Hemiptera (Grubbs et al., 2015; Khan et al., 2017; Vargas-Lowman et al., 2019). In the five species we tested (**Figures 1, 3**) and consistently with previous reports in *Papilio, Bicyclus*, and *Bombyx* (Liu et al., 2021; Tatematsu et al., 2011), *scarlet*-mutant eyes showed residual yellow or reddish pigmentation compared to *white*-mutants eyes. These light-color pigments may be pteridines, based on a previous report of ekapterin in *Helicoverpa* moths (Khan et al., 2017), but this hypothesis will require further investigation in more species. Overall, while the nature of light-colored pigments remains relatively elusive, it is clear that *scarlet*-dependent ommochromes are responsible for the dark pigmentation of adult eyes in many lepidopteran lineages and beyond.

**Figure 7.**
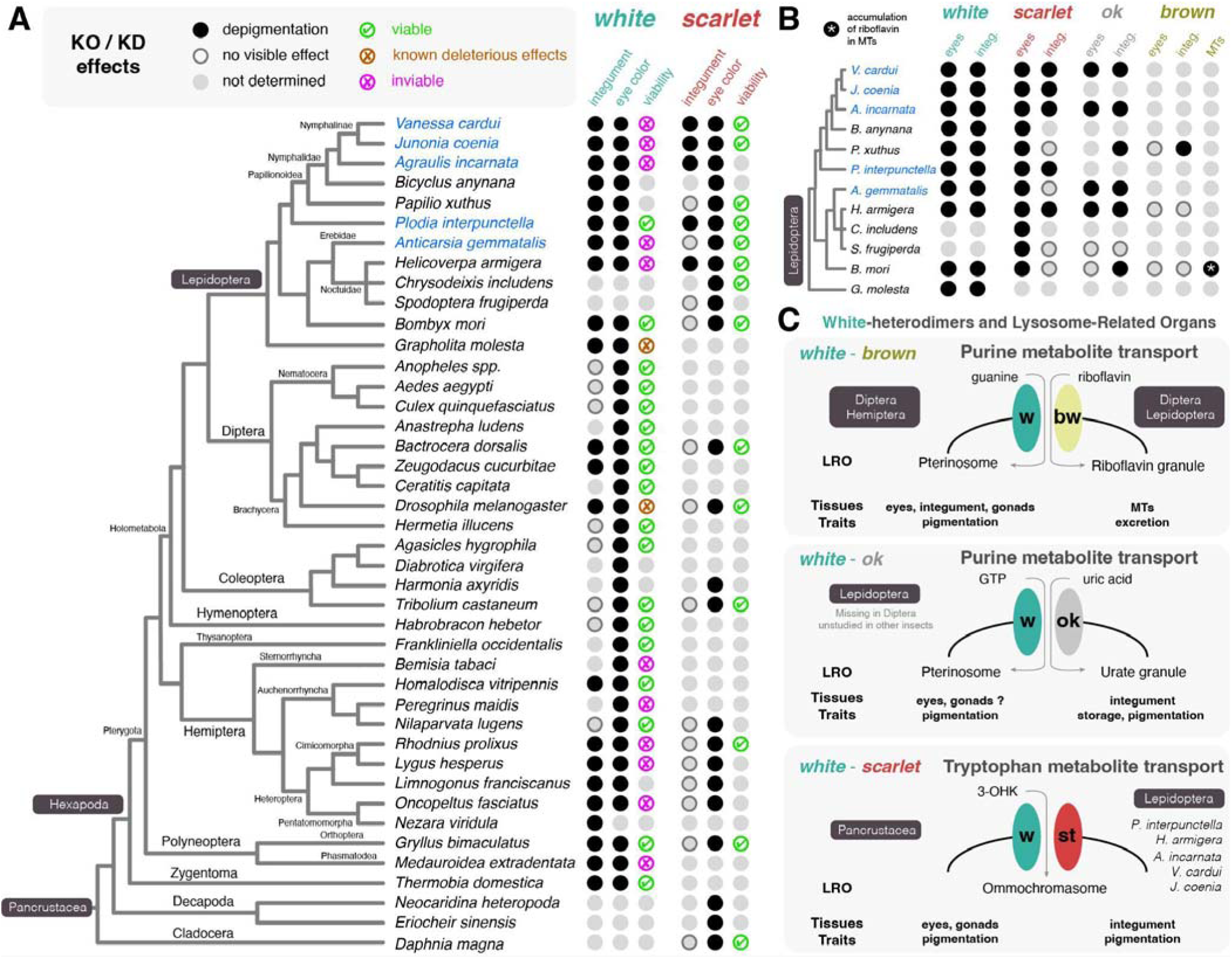
Summary of ABCG transporter gene loss-of-function phenotypes across insects and crustaceans. **A.** Literature review of *white* and *scarlet* perturbation studies across 41 pancrustacean species (full dataset and list of references in **Supporting Information Table S4**). KO: gene knockouts (spontaneous or CRISPR-based). KD: RNAi knockdowns. Magenta: species in which *white* effects could be documented in mosaic clones or in transient RNAi phenotypes, but could not generate mature adults, due to strongly deleterious or recessive-lethal effects. Black dots indicate depigmentation phenotypes at any stage. Grey circles indicate that there were no conspicuous coloration phenotypes other than in ocular structures. Green check-marks: mutants could be bred to the homozygous state without visible deleterious effects. Magenta crosses: homozygous germline mutants are inviable. Orange crosses: homozygous *white* mutants reach the adult stage but deleterious effects on size, longevity or behavior are documented. **B.** Summary of perturbation studies in Lepidoptera with available data for *brown* and *ok*. MTs: Malpighian Tubules. **C.** Model for the differential pleiotropy of *white* and *scarlet* mutants in Lepidoptera. White functions both in the transport of purine derivatives and 3-OHK, whereas Scarlet functions in a smaller set of less essential tissues. Some of the integumentary effects of *scarlet* KOs in nymphalid butterflies are not fully understood and require further investigation.

At deeper phylogenetic levels, CRISPR KOs of *scarlet* also result in eye depigmentation in the *Daphnia* water fleas and in decapod crustaceans (Fatimah et al., 2022; Ismail et al., 2018; Li et al., 2022; Maar et al., 2024). While *white* KOs have not been attempted in these studies of crustaceans due to the existence of multiple *white* paralogs, they yield white-eyed phenotypes in other insect lineages including in wasps, crickets, stick insects, and silverfish (Bai et al., 2024; Di Cristina et al., 2025; Gonzalez-Sqalli et al., 2024; Inada et al., 2025; Matsuoka & Monteiro, 2018; Ohde et al., 2018). From this phylogenetic framework, we infer that the White-Scarlet heterodimer functions as a conserved transporter of ommochrome precursors functioning in eye pigmentation across Pancrustacea. It remains to be tested whether this role is conserved in other arthropods such as spiders, where the expression of candidate ABCG transporter orthologs and ommochrome pathway enzymes have been described (Croucher et al., 2013).

### Mutations of *scarlet* show less deleterious pleiotropy than for *white*

Our inability to obtain non-mosaic (G_1_) homozygotes highlights the deleterious nature of *white* mutations in two butterfly species (*V. cardui* and *J. coenia*). Similar inviability effects have been described in other species of Lepidoptera (Khan et al., 2017; Su et al., 2022), Hemiptera (Heu et al., 2022; Klobasa et al., 2021; Lima et al., 2024; Reding & Pick, 2020), and Phasmatodea (Di Cristina et al., 2025), while *white* mutants are viable in *D. melanogaster* but sterile in *Drosophila suzukii* (Yan et al., 2020). It is worth considering a few possible mechanisms that may explain these lethal effects for future investigation. Recent studies in *D. melanogaster* showed a direct impact of *white*-deficiency on locomotor activity, egg-laying behavior, and overall gene expression (Ferreiro et al., 2018; Rickle, Sudhakar, et al., 2025), and at least some of these effects can be tied to reduction in biogenic amine levels such as serotonin, dopamine, and histamine (Borycz et al., 2008; Kain et al., 2012; Sitaraman et al., 2008). In addition, White interacts with Brown, as well as its paralog Ok in Lepidoptera, to transport purine compounds in Malpighian Tubules **(Figure 7C)**. Here as well, there is emerging evidence that perturbation of this transporter system could have broad systemic effects. First, in addition to their roles in pterin pigmentation, White-Brown heterodimers are important for the formation of lysosome-related organelles (LROs) that accumulate riboflavin in the renal Malpighian Tubules (MTs) of *Drosophila* and *Bombyx* (Van Breugel, 1987; Zhang et al., 2018). Proper riboflavin storage in the tubules might regulate the systemic supply of this essential nutrient (vitamin B2), which plays critical roles in mitochondrial function and insect physiology (Manole et al., 2017). Second, many insects excrete nitrogen in the form of purine derivatives such as uric acid (Weihrauch & O’Donnell, 2021), and the White-Brown dimer (or the White-Ok dimer in Lepidoptera) likely supports this via purine uptake by the MTs, with Brown-White required for riboflavin LROs and potentially buffering purine metabolism and waste handling (Sullivan et al., 1979; Zhang et al., 2018). In a nutshell, the multiple functions of White extend beyond coloration and could explain its lethality in some of the tested species (**Figure 7A**). It remains unclear whether roles in neurotransmission, nitrogenous waste excretion and storage, regulation of riboflavin levels (Rickle, Krittika, et al., 2025), or other functions explain the molting defects we observed in *V. cardui* and *J. coenia* and *white* mutants.

In contrast, Scarlet functions more narrowly in ommochrome transport, and its loss-of-function alleles are generally viable across taxa, with eye coloration changes uncoupled from systemic lethality. This pattern highlights the asymmetric pleiotropy of ABC transporter dimeric partners (**Figure 7C**), where White is highly pleiotropic and Scarlet comparatively specialized.

### Complementary roles of White-Scarlet and White-Ok in caterpillar coloration

Relatively little is known about the molecular basis of coloration in lepidopteran caterpillars, particularly in contrast to the extensive work on adult wing pigmentation. The larval epidermis consists of a single cell layer that secretes the cuticle. While melanins deposited in the cuticle can provide black pigmentation, epidermal cells can also accumulate other classes of pigments. Our CRISPR knockout analyses indicate that White heterodimers have specialized roles in the accumulation of two distinct pigment classes, urate and ommochrome.

Urate granules are LROs that provide an opaque, white coloration to larval epidermal cells and have been described in several lepidopteran families (Buckner & Newman, 1990; Fujii & Banno, 2019; J. Lee et al., 2018; Lhonoré et al., 1980; Ninomiya et al., 2006). The White-Ok heterodimer is required for the formation of urate granules in *Bombyx* and *Helicoverpa* caterpillars (Khan et al., 2017; Kômoto et al., 2009). Consistent with this role, we show that null mutations of *white* and *ok* result in a translucent larval integument in *V. cardui*, as well as loss of white pattern elements in *A. incarnata*, *J. coenia*, and *A. gemmatalis*. Similar translucency effects were also observed in pupae. Together, these results indicate that urate granules are commonly used not only to generate white coloration but also to confer opacity to the integument of lepidopteran larvae and pupae. In addition, the integument discoloration phenotypes observed in *white* and *scarlet* knockouts; particularly pronounced in early larval instars of *V. cardui* and *J. coenia*, consistently visible across all larval stages of *A. incarnata*, and most evident in late larval stages of *P. interpunctella;* indicate that orange, red, and brown pattern elements in these species are based on ommochrome pigmentation.

Interestingly, we did not detect any effect of *white*, *scarlet*, and *ok* knockouts on wing coloration in any of our tested species. These negative results further anchored the idea that scale ommochrome pigmentation uses alternative precursors and transporters (How et al., 2023; Uchiyama et al., 2023). Whether *ok* has a role in pteridine scale pigmentation could be assessed in Pieridae, a lineage of butterflies whose scale coloration is mostly based on pteridines (Descimon, 1975; Wijnen et al., 2007).

### Implications for marker development

These comparative data help explain why *scarlet* is emerging as a particularly useful marker gene for genome editing in Lepidoptera and beyond. The viability and non-deleterious nature of *scarlet* mutations make them advantageous for CRISPR-Cas9 screening, both as direct eye-color reporters and as co-CRISPR markers for mosaic efficiency. In contrast, *white* mutations, although highly penetrant in pigmentation, carry a higher risk of confounding pleiotropic effects on viability or fertility, thereby limiting their general utility. This is particularly true in Lepidoptera and Hemiptera.

Conversely, *scarlet* mutations often produce visible eye discoloration phenotypes across insects and even crustaceans (**Figure 7A**). We recommend the targeting of *scarlet* in lieu of *white* for implementing the co-CRISPR strategy (Kane et al., 2017): when targeting a gene of interest, co-targeting *scarlet* should generate mosaic eye phenotypes that mark individuals in which CRISPR reagents were successfully delivered. Eye depigmentation can also facilitate the screening of fluorescent reporters such as *3xP3*-driven fluorescent proteins, and we recommend the use of *scarlet* knockout lines, instead of *white* mutants (Berghammer et al., 1999; Heryanto, Mazo-Vargas, et al., 2022; Shodja et al., 2025), for establishing transgenic stocks in emerging model insects in conjunction with eye-specific transgenesis markers (Kane et al., 2017). It is also worth noting that the yellow pigmentation of male larval testes in *P. interpunctella scarlet* mutants (**Supporting Information Fig. S4**) can facilitate sorting transgenic founders of a given sex when establishing transgenic strains, whereas *white* mutants lack this feature and require sex determination at the pupal stage (Heryanto, Hanly, et al., 2022). Last, the depigmentation and translucency effects of *scarlet* KOs we observed on lepidopteran caterpillars and pupae can facilitate the detection of other *in vivo* fluorescent marker, directly through the cuticle, such as the glial activity of *3xP3*, the broad activity of viral promoters such as *hr5/ie1*, or the activity of the *FibL* promoter in silk glands (Shodja et al., 2025). In summary, although *white* mutants have historically been a genetic tool of choice for functional genetics in insects, *scarlet* mutants can be equally practical, while also avoiding potential deleterious effects.

## Acknowledgments

We are very grateful to Kitty Brian for her continued love and care in rearing butterflies and for sharing her *Agraulis incarnata* stock. We also thank Martina Tsimba for assistance with caterpillar rearing. We are thankful to Rachel Canalichio of the GWU Harlan Greenhouse, as well as Jorge Fidhel Gonzalez and Amy Eddings at the Duke Biology Research Greenhouses, and their teams, for their support in growing host plants.

## Data Availability Statement

The data underlying this article are available in the article and in its online Supporting Information. https://doi.org/10.6084/m9.figshare.31395594

## Supporting Information

**Supplementary Figure 1.**
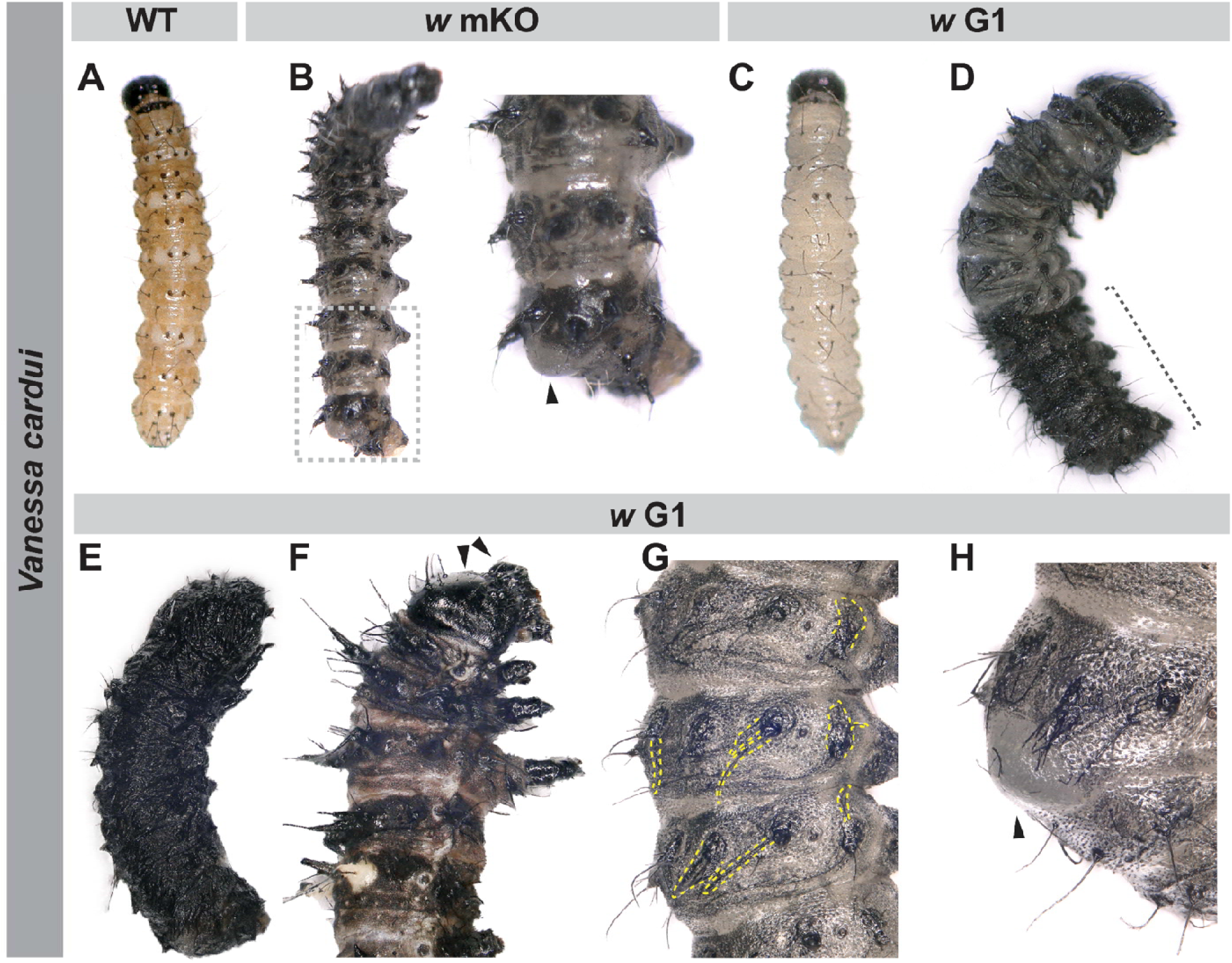
CRISPR-Cas9 disruption of the *white* gene in *V. cardui* causes severe defects at early larval stages. **(A)** WT L1 caterpillar **(B)** Third instar G_0_ *w* crispant with bristle defects and posterior bulge (black arrowhead) **(C-H) Range of defective phenotypes in G_1_ *V. cardui white* mutants. (C)** An L1 *w* G_1_ individual with complete loss of integument pigmentation (compare with A) **(D)** L2 *w* G_1_ individual with molting defects. Dotted bracket marks the posterior segments of the caterpillar with old, unshed integuments. **(E)** L2 *w* G_1_ fully encapsulated in its old integument **(F)** L2 *w* G_1_ with severe integumentary defects and retention of the previous head capsule following an incomplete larval molt (black arrowheads) **(G)** L2 *w* G_1_ with double cuticle appearance due to incomplete molting. New bristles appear under the old, unshed integument (a portion of them denoted by yellow dotted outlines). **(H)** the posterior bulge (black arrowhead) of the same individual in panel G.

**Supplementary Figure 2.**
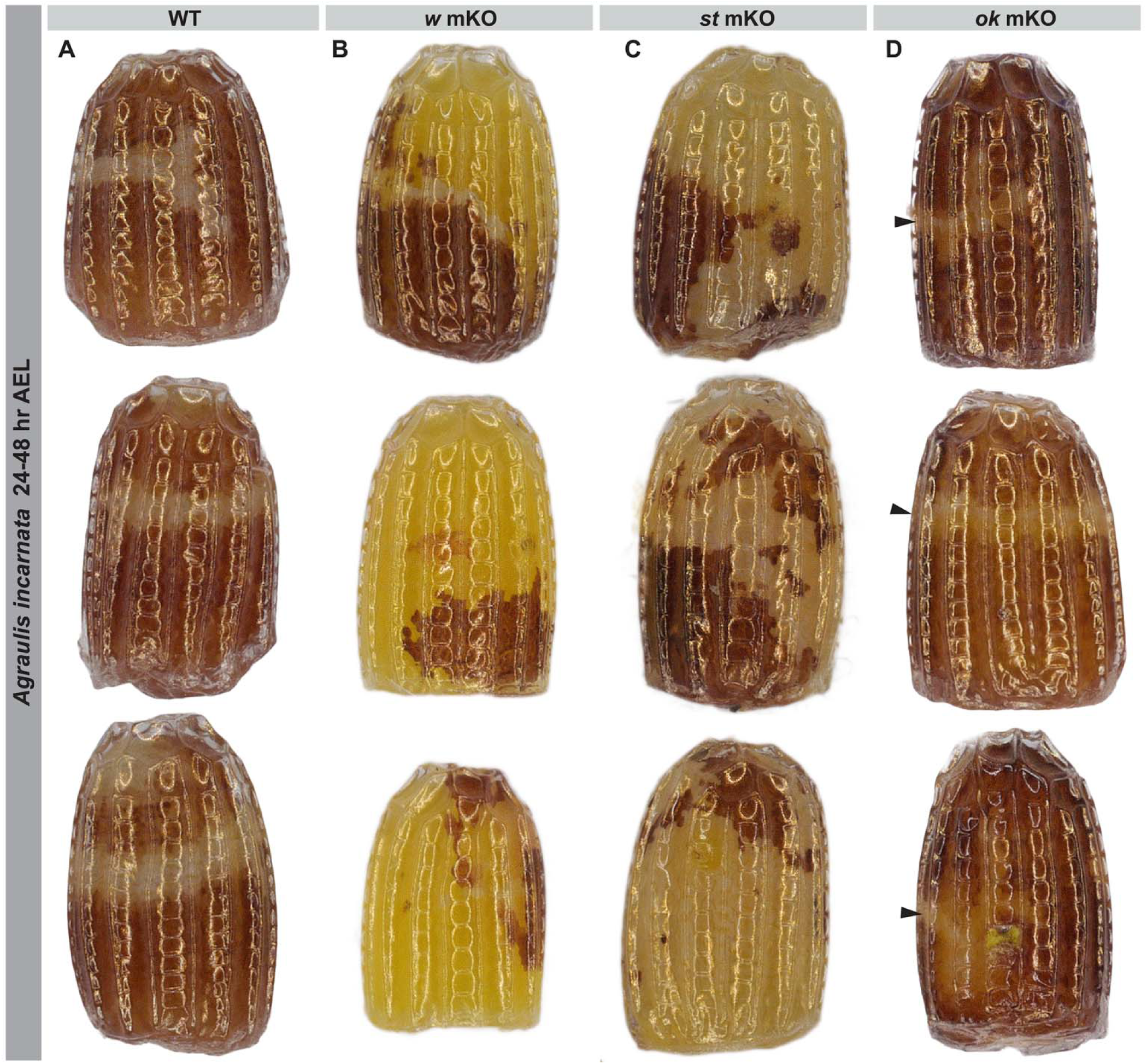
Pigmentation of the egg serosa in *Agraulis incarnata* WT, *white* mKOs, *ok* mKOs, and *scarlet* mKOs embryos. All eggs were imaged between a 24-48 hours AEL (after egg laying). Three representative phenotypes are depicted for each genotype. **(A)** WT eggs display a brownish-red/dark orange pigment in the serosa, in addition to a white fringe. (**B**) *white* mKO eggs lose both dark and white pigmentation, presenting a vivid yellow pigmentation. Similar effects are observed in the (**C**) *scarlet* mKO developing eggs. (**D**) In *ok* mKO eggs, the thin white fringe regions changed to yellow (black arrowheads).

**Supplementary Figure 3.**
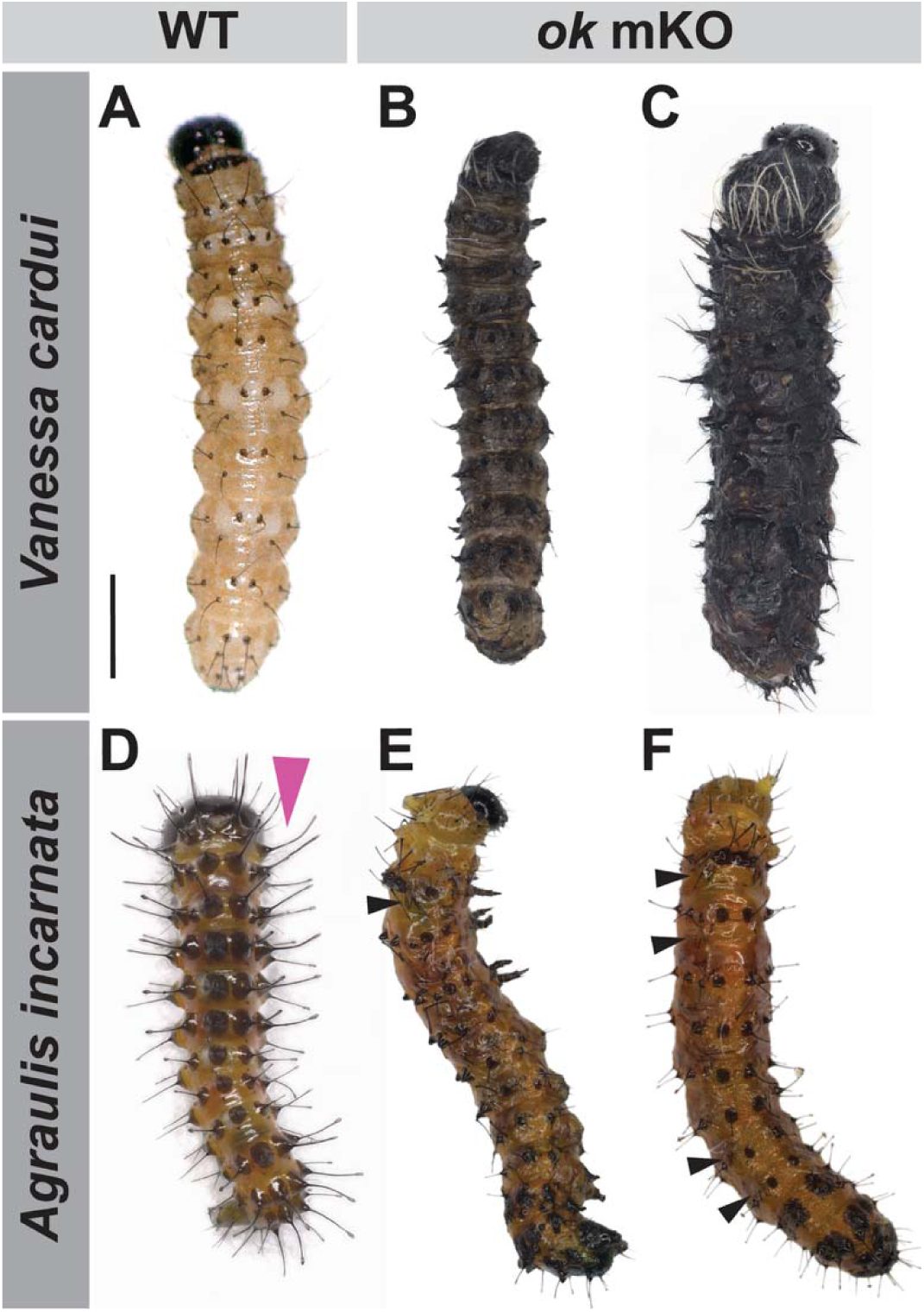
CRISPR-Cas9 disruption of the *ok* gene in *V. cardui* and *A. incarnata* causes molting defects at early larval stages. (A-C) *V. cardui* G_0_ individuals with molting defects appear darker with thickened integument and defective bristles (compare B and C to A). **(D-F)** Examples of *A. incarnata ok* mKO G_0_ individuals that have failed to molt and exhibit a double integumentary phenotype. Black arrowheads indicate bristles entrapped in old, unshed integument. (compare E andF to D).

**Supplementary Figure 4.**
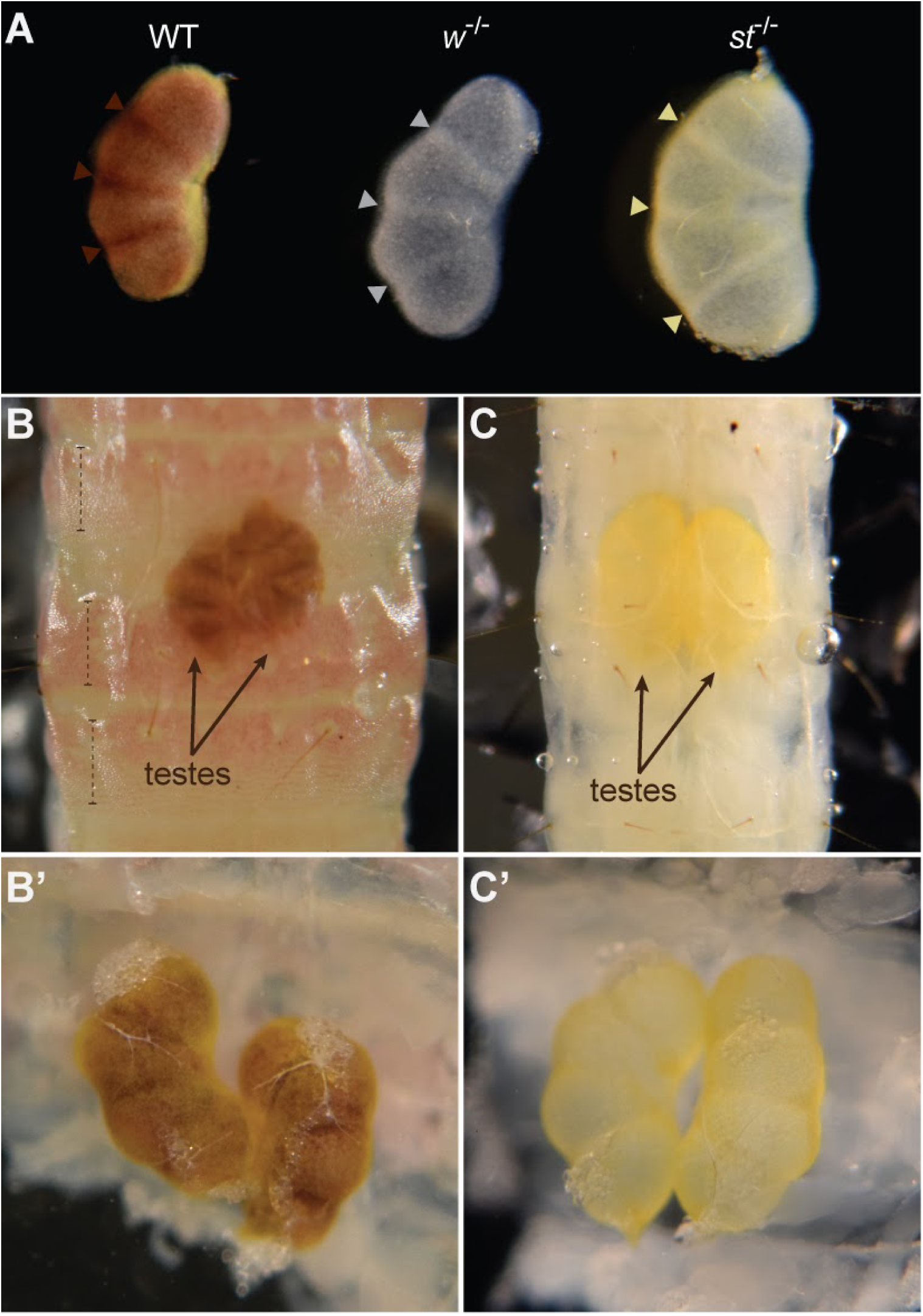
Phenotype of *Plodia interpunctella* fifth instar larval testes: *scarlet* mutants can be used to distinguish males and females. **(A)** Comparison of dissected testes from wild type (left), *w^-/-^* (middle), and *st^-/-^* (right) fifth instar larvae. In WT larvae, brown testicular masses are conspicuous on the dorsal abdomen and can be used for distinguishing males from females, with three dark reddish brown stripes of septa (arrowheads) that connect 4 sac-like follicles (Ashrafi et al., 1972). The wFog strain (*w^-/-^*) testes are unpigmented, while *st^-/-^*testes are yellow. **(B)** Dark brown kidney shaped testes (arrows) in the mid-dorsal abdomen are visible through the cuticle of an early fifth instar wild type larva. Dashed lines point to the pink pigmentation on the larval body wall. **(C)** Yellow testes are visible through the transparent larval cuticle of ta *st^-/-^* mutant fifth instar. The pair of wild type **(B’)** and *st^-/-^* **(C’)** testes from panels B and C, respectively, partially dissected out but connected to their larval cuticles.

**Figure S5.**
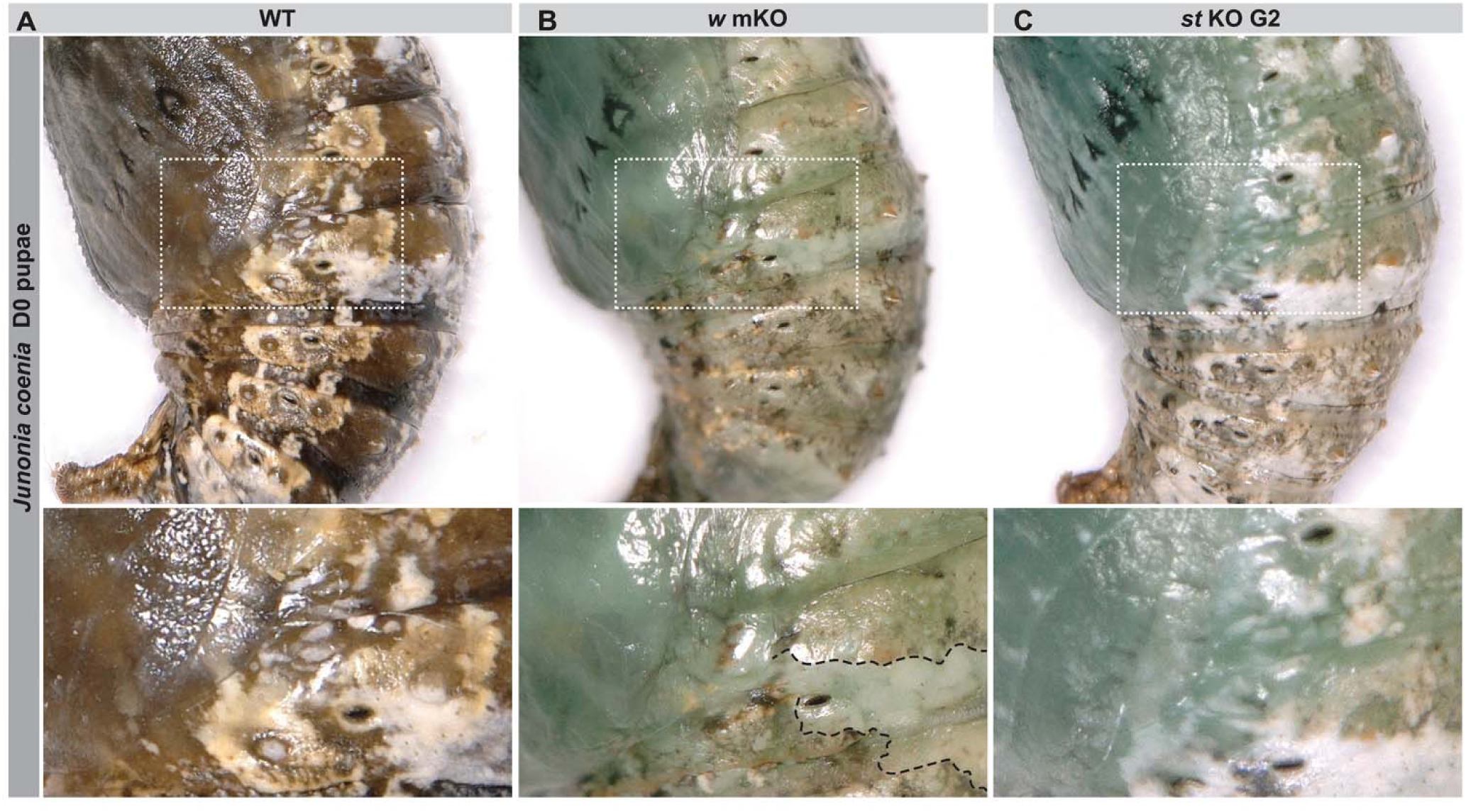
Effects of *white* and *scarlet* knockout on the opacity of pupal integuments in *J. coenia*. Lateral view of 0–4 hr APF pupae (ventral side to the left), showing posterior segments. **(A)** WT pupae are green-brown with white colored patterns along the lateral side. **(B)** *white* mKO and **(C)** *scarlet* KO individuals appear light blue-green. White-colored patches are disrupted in *white* mKO mutants (dotted lines in B, inset) but not in *scarlet* KO mutants.

**Supplementary Figure 6.**
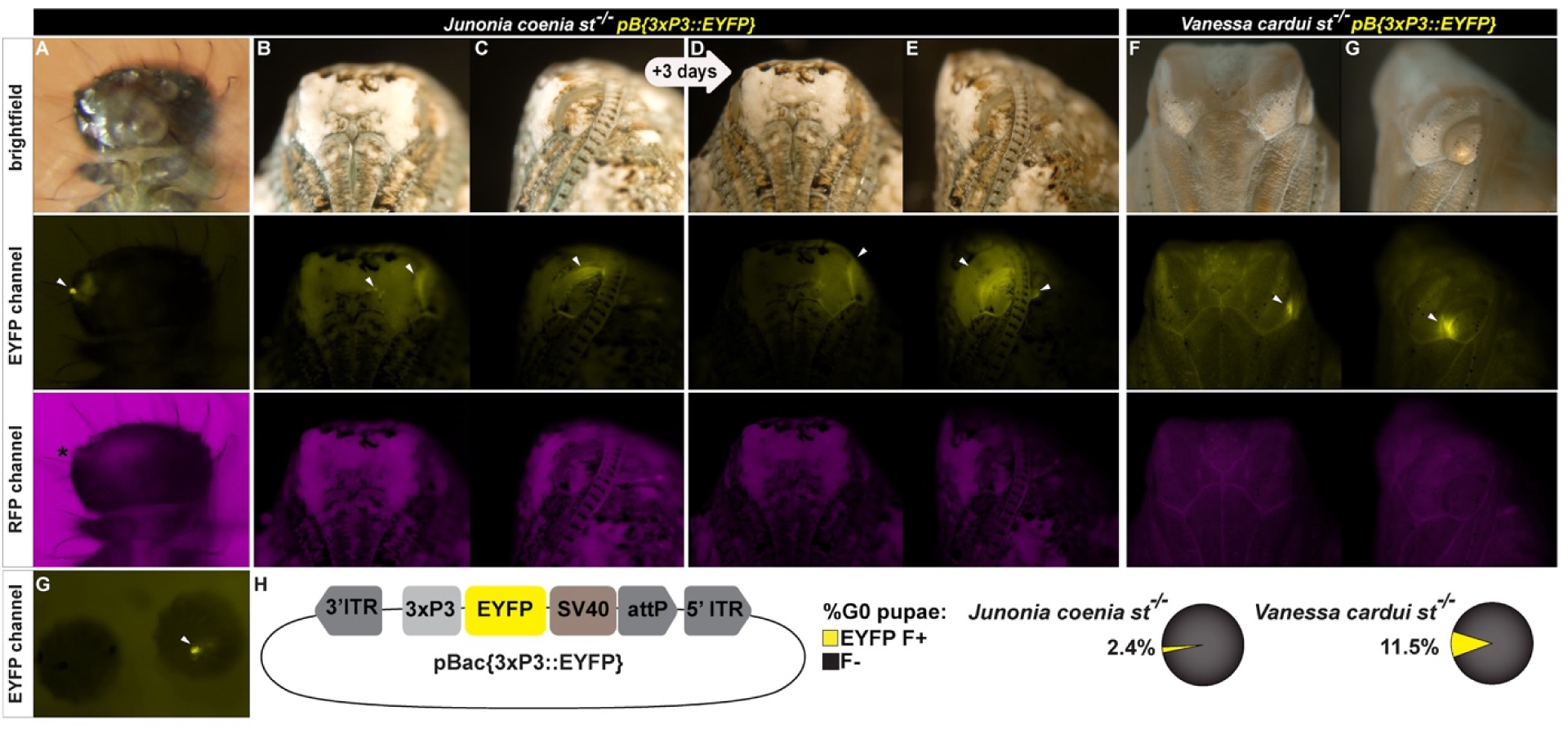
Transgenic mosaic *J. coenia* and *V. cardui* scarlet mutants carrying the *[3xP3::EYFP]* expression cassette following pBac mediated integration. Each G0 individual is shown through a brightfield channel (top), EYFP channel (middle), and RFP channel (bottom). **(A)** Ventral view of unilateral ocelli expression (arrowhead) in a G0 *J. coenia* hatchling. **(B)** ventral and **(C)** lateral views of mosaic *[3xP3::EYFP]* detection in the developing eye at an untimed pupal stage, and **(D, E)** stronger expression in the same pupa after 3 days. **(F, G)** Example of mosaic *[3xP3::EYFP]* expression in a developing *V. cardui st*^-/-^. (**H**) schematic of the pBac*[3xP3::EYFP]* donor plasmid and the proportion of fluorescent positive G_0_ individuals detected in each species.

**Supplementary Figure 7.**
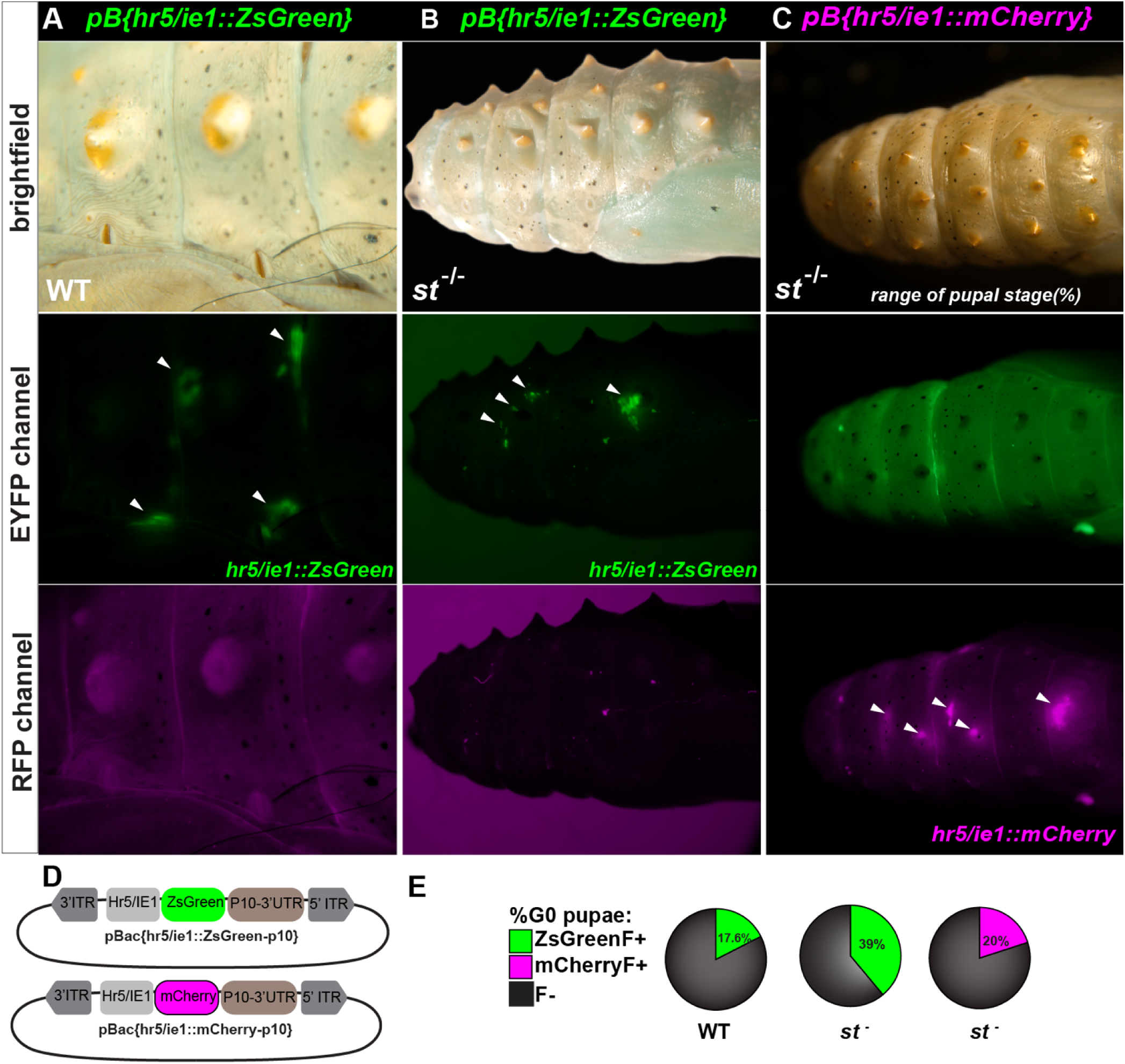
Mosaic expression of the hr5/ie1 promoter in *V. cardui* wild-type and *st^-/-^* mutant backgrounds, an alternative marker to eye/glia. Two different donor plasmids were tested to drive green and red fluorescent proteins with the same promoter: (**A**) *pB[hr5/ie1::ZsGreen]* in a WT background, (**B**) *pB[hr5/ie1::ZsGreen]* in a *scarlet* mutant line background, and (**C**) *pB[hr5/ie1::mCherry]* in a *scarlet* mutant line. These were assayed across 1, 2, and 1 injection trial(s), respectively. Each represented G0 individual is shown through a brightfield channel (top), EYFP channel (middle), and RFP channel (bottom). **(A)** Lateral view of a WT pupa injected with *pB[hr5/ie1::ZsGreen]* displays localized patches of fluorescence in a spatially random pattern. (**B**) Lateral view of a *scarlet* mutant pupa injected with *pB[hr5/ie1::ZsGreen]* displays brighter, easily detectable patches of fluorescence compared to the WT injected. The RFP channel serves as a control for autofluorescence and artifacts. (**C**) Dorsal view of a *scarlet* mutant pupa injected with *pB[hr5/ie1::mCherry]* showing red fluorescent signal, absent in the EYFP channel indicating true fluorescence. (**D**) schematics of each donor plasmid: *hr5* (*homologous region 5*) functions as an enhancer element of the *ie1* promoter. (**E**) Proportion of fluorescent positive G_0_ individuals detected for each plasmid. In a *scarlet* mutant background, the proportion of positive G_0_ pupae increased to 39% (37 of 94 pupae across two trials) (**Supporting information Table 3**). Detection of fluorescence was markedly easier in the *st^-/-^* mutants fluorescent patches in each individual were more abundant, spatially larger, and qualitatively brighter compared with those observed in the WT.

**Supplementary Figure 8.**
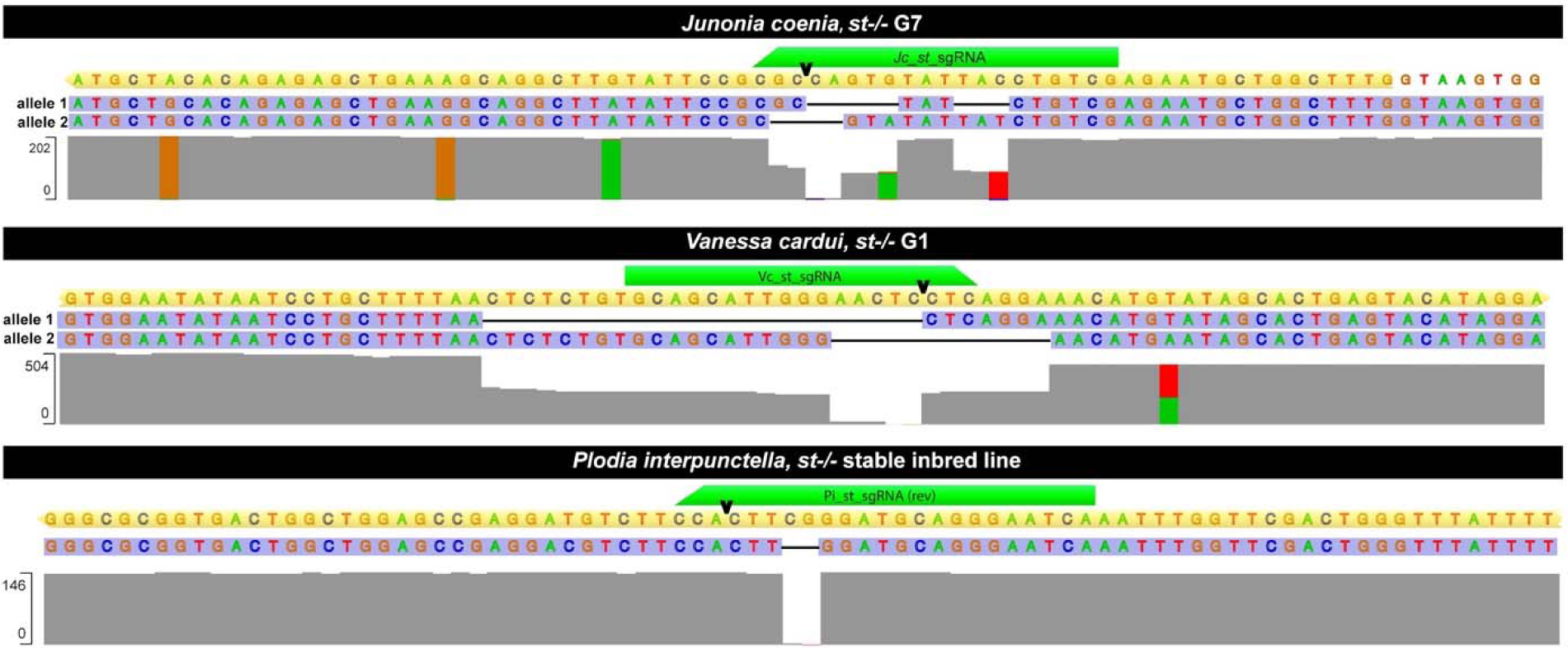
Genotyping of stable *scarlet* mutant lines of *J. coenia*, *V. cardui*, and *P. interpunctella*. In each panel, the top sequence represents the reference amplicon (WT) where yellow indicates the coding region and directionality. Deletions in mutant alleles are represented with missing sequence alignment. The grey coverage plot shows read depth across the reference sequence. The number of aligned reads (y-axis) at each nucleotide position (x-axis) illustrates sequencing depth and uniformity across the reference amplicon. Regions of low coverage correspond to deletions in the scarlet mutant alleles.The mutant alleles are: *J. coenia* (allele1: c.1466_1476delinsTAT, allele2: c.1464_1467delGCCA), *V. cardui* ( allele1: c.1420_1443delCTCTCTGTGCAGCATTGGGAACTC, allele2: c.1413_1424delAACTCCTCAGGA), and *P. interpunctella* (allele: c.171_172delCG).

